# Recurrent oncogenic ZC3H18 mutations stabilize endogenous retroviral RNA

**DOI:** 10.1101/2025.01.10.632423

**Authors:** Tanzina Tanu, Anna M. Cox, Jennifer Karlow, Priyanka Sharma, Xueyang He, Constance Wu, Swathy Babu, Jared Brown, Kevin M. Brown, Stephen J. Chanock, David Liu, Tongwu Zhang, Kathleen H. Burns, Paul L. Boutz, Megan L. Insco

## Abstract

Endogenous retroviral (ERV) RNA is highly expressed in cancer, although the molecular causes and consequences remain unknown. We found that ZC3H18 (Z18), a component of multiple nuclear RNA surveillance complexes, has recurrent truncating mutations in cancer. We show that Z18^trunc^ mutations are oncogenic and that Z18 plays an evolutionarily conserved role in nuclear RNA surveillance of ERV RNA. In zebrafish, Z18^trunc^ expedited melanoma onset and promoted a specific accumulation of ERV RNA. Z18 mutant human cell lines from the Cancer Cell Line Encyclopedia also expressed higher levels of ERV RNA. In engineered human melanoma cells, Z18^trunc^ enhanced ERV RNA accumulation more than loss of one Z18 copy, indicating dominant negative activity. Z18^trunc^ directly bound and stabilized ERV RNA. Notably, expression of ERV RNA was sufficient to expedite oncogenesis in a zebrafish model, which is the first evidence of which we are aware that ERV transcripts can play a functional role in cancer. Our work illuminates a mechanism for elevated ERV transcripts in cancer and supports that aberrant RNA accumulation is broadly oncogenic.

## Main Text

Healthy cells require appropriate levels of high-quality RNA, therefore RNA dysregulation contributes to many diseases, including cancer. RNA homeostasis is regulated through RNA production and destruction. While the role of RNA production through transcription factors, chromatin regulation, and enhancers in cancer has been well studied (*1*), less is understood about how RNA degradation pathway perturbation contributes to cancer. Mechanisms that detect and degrade aberrant or unstable RNAs in the nucleus have only recently been described (*2–5*) and we previously discovered that deficient nuclear RNA surveillance of aberrant prematurely terminated RNAs (ptRNAs) is oncogenic (*6*). Nuclear RNA surveillance components are mutated in up to 21% of melanomas, and recurrent mutations occur in two components of the complex that clears ptRNAs, including ZC3H18 (*6*) (hereafter referred to as Z18), suggesting that deficient nuclear RNA surveillance is a widespread contributor to human tumorigenesis.

Different types of unstable or aberrant RNA in the nucleus are recognized for degradation by different protein complexes. The PolyA Exosome Targeting (PAXT) complex identifies aberrant polyadenylated transcripts (*4*), including ptRNAs (*7*) that we previously found to be oncogenic (*6*). The Nuclear Exosome Targeting (NEXT) (*3*) complex identifies a surprising range of non-coding RNAs for degradation, including promoter upstream transcripts (PROMPTs), enhancer RNAs (eRNAs) (*8*), long interspersed element-1 (LINE-1) retrotransposons (*9*), and long terminal repeat (LTR) retrotransposons including endogenous retroviral (ERV) RNAs (*10*). Retrotransposons are repetitive genomic sequences that propagate via an RNA intermediate and include LINEs, short interspersed elements (SINEs), and retroviral-derived ERVs and LTRs. ERVs and LTRs make up about 8% of the human genome (*11*) with both having accumulated mutations that render them non-infectious and immobile (*12*). However, ERV sequences may retain enhancer activity (*13*), Pol II promoter activity (*14*), splice sites, and polyadenylation sequences, which can contribute to the production of long noncoding RNAs and chimeric transcripts as well as fragmented open reading frames (ORFs) with protein coding potential (*15*). Transposable elements have been reported to be expressed at increased levels in multiple cancer types, with the ERV class of retroelements being most affected (*16, 17*). ERV upregulation in cancer has been shown to correlate with increased immune infiltration and immune therapy efficacy in multiple cancers (*17–20*). However, the causes and cancer cell intrinsic functional consequences of ERV upregulation in cancer are unknown. Whether elevated levels of ERV transcripts or proteins are pathogenic or an epiphenomenon of broader transcriptional changes in malignant tissue remains unclear.

In this study, we show that recurrent truncating mutations in Z18 produce a truncated protein which retains its ability to bind RNA and loses the domain required to recruit RNA degradation machinery. The truncated isoform accelerates melanoma onset in a zebrafish model and promotes a specific accumulation of ERV RNAs without affecting the steady-state levels of other aberrant/unstable nuclear RNAs. We find that the role of Z18 in ERV degradation is conserved across species and that Z18 truncating mutations function in a dominant negative manner both in zebrafish and human melanoma cells. The truncated isoform binds ERV RNA, protecting it from nuclear degradation. Importantly, expression of an ERV was sufficient to expedite melanoma onset in a zebrafish melanoma model. By studying Z18 mutations in melanoma, we have revealed a functional role for ERV RNA in oncogenesis. This study expands the idea that accumulated aberrant RNA contributes to cancer phenotypes.

## Results

### Recurrent ZC3H18 Truncating Mutations are Oncogenic

We previously found that CDK13 activates the PAXT complex to target aberrant ptRNAs for degradation. When CDK13 is mutated, ptRNAs are stabilized, leading to more aggressive melanoma (*6*). As Z18 is a component of PAXT, is understudied, and because we previously found Z18 was a low frequency driver in melanoma using OncodriveFM (*6*), we further investigated Z18. Using The Cancer Genome Atlas (TCGA) melanoma (*21*) patient samples, PolyPhen-2 (*22*) was used to predict the impact of mutations on protein structure/function and found that melanoma had an enrichment of Z18 deleterious mutations (P=0.0047). In a larger cohort of cutaneous melanoma patients (n = 1347), we observed that 35% of patients with available copy number data had biallelic or monoallelic loss of Z18 (291/823). 78% of Z18 mutations were deleterious by PolyPhen-2, 50 patients had deleterious mutations (7 truncating and 43 deleterious missense) out of 64 patients with Z18 mutations (Data S1, “Z18 Mut & Melanoma Patient Refs”). Other TCGA cancers also showed significant enrichment of deleterious mutations from PolyPhen-2 including: uterine corpus endometrial carcinoma (P=0.0041), cervical squamous cell carcinoma (P=0.028), colon adenocarcinoma (P=0.013), and lung adenocarcinoma (P=0.016) (Data S1, “PolyPhen-2 Results”). These data suggest that Z18 deleterious mutations and copy number loss are selected for in melanoma and potentially in other cancers.

We then investigated Z18 mutations in a larger cohort of publicly available data (Data S1, “Pan-cancer Patient Sample Refs”). We identified 506 deleterious Z18 mutations from 23,411 patients and found that truncating mutations were statistically enriched in the region preceding the RNA surveillance binding domain (aa 679-901) as compared to the rest of the protein (odds ratio=5.89, n=23411, P=1.66e-17, Fisher’s exact test) (Fig. 1A, Data S1, “Z18^trunc^ Fisher Exact Test” and “Pan-cancer Z18 Mutations”). Specifically, 49 patients had a truncating mutation at R680 and 37 patients had truncating mutations just downstream, suggesting that the loss of the distal Z18 C-terminal domain is oncogenic. Z18 truncating (Z18^trunc^) mutations were most frequently found in endometrial carcinoma (20/67), stomach adenocarcinoma (18/29), and colorectal adenocarcinoma (14/35), which are tumors that can have microsatellite instability (MSI-high), so we asked whether Z18 mutations were more likely to occur in MSI-high tumors. In TCGA patients, Z18 mutations were more likely to occur in MSI-high tumors (24%, 75/319 MSI-high vs. 1%, 137/10463 microsatellite stable); although most Z18 mutations occurred in microsatellite stable tumors (65%, 137/212) (Table S1). As loss-of-function mutations are predicted to be spread evenly across a gene, enrichment of Z18 truncating mutations suggests an additional function such as dominant negative or neomorphic activity. The Z18 C-terminal domain is required to recruit RNA surveillance machinery (*23*), thus we hypothesized that these mutations would result in an oncogenic isoform that fails to degrade target RNA, contributing to oncogenesis.

**Figure 1.**
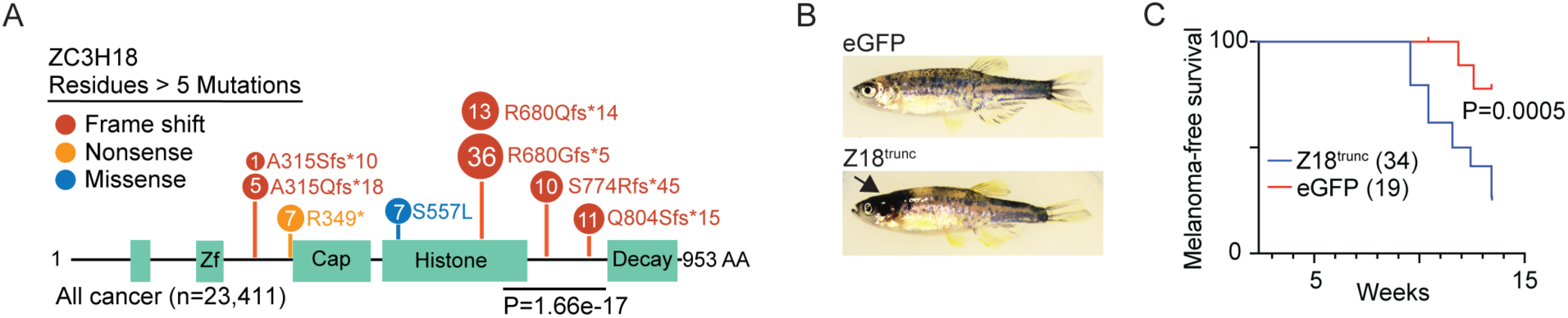
Recurrent ZC3H18 Truncating Mutations are Oncogenic. A) Z18 lollipop plot from patients (pan-cancer). P=1.66e-17 (Fisher’s exact test comparing truncating mutations in residues 679-901 vs. the rest of the protein). n= patients. B-C) Triples zebrafish with melanocyte-specific expression of eGFP or human Z18^trunc^ (R680Gfs*5). B) Representative photos at 9 weeks. Arrow indicates early melanoma. C) Percent melanoma-free survival. P=0.0005 (log rank). n= zebrafish.

To test whether Z18^trunc^ is oncogenic in melanoma, we used the zebrafish MAZERATI(*24*) rapid genetic modeling system which allows melanocyte-specific gene expression in first generation animals. The MAZERATI system was used in *BRAF^V600E^; p53-/-*; *mitfa-/-* zebrafish, subsequently referred to as the ‘Triples’. In addition to having human oncogenic BRAF expression and *p53* mutation, these fish lack Mitfa, the master regulator of the melanocyte lineage, and thus lack melanocytes (*24–26*). Single-cell embryos were injected with vectors expressing Mitfa, thus rescuing melanocytes and expressing genes of interest. As we previously demonstrated that ptRNA degradation mechanisms were evolutionarily conserved between humans and zebrafish (*6*) and human and zebrafish Z18 are highly similar (67.62% nucleotide sequence and 62.9% protein sequence by BLAST (*27*)), we tested the function of the most frequent patient truncating mutation, Z18 R680Gfs*5 (hereafter Z18^trunc^), in this zebrafish melanoma model (Fig. S1A). Z18^trunc^ expression produced increased black patches at 9 weeks (Fig. 1B, Fig. S1B) and expedited melanoma onset compared to eGFP-expressing control (Fig. 1C, S1D, biologic replicate in S1C). These data show that human Z18^trunc^ expression expedites melanoma onset in zebrafish.

### ZC3H18^trunc^ Promotes ERV Accumulation

As Z18 is a component of multiple RNA surveillance complexes, we investigated whether RNA surveillance substrate RNAs were stabilized by Z18^trunc^ expression. Since we previously observed Z18 bound to the PAXT complex (*4*) that clears polyadenylated ptRNAs (*6*), ptRNAs were quantified from pA-selected bulk RNA-sequencing from zebrafish tumors with melanocyte-specific expression of eGFP (n=3) or Z18^trunc^ (n=3) (Fig. S2A). We found no significant upregulation of CDK13-dependent ptRNAs in the Z18^trunc^ zebrafish melanomas as compared to controls (Fig. S2B). As Z18 is also reported to be a component of the NEXT nuclear RNA surveillance complex (*3*), we assayed NEXT substrate RNAs. The NEXT complex was recently reported to degrade LINEs (*9*), whose transcripts are typically polyadenylated. Transposable element (TE) expression from pA-selected RNA-seq was measured with SQuIRE, which uses stringent mapping coupled with an expectation-maximization based algorithm (*28*). Average TE expression was calculated for eGFP (n=3) and Z18^trunc^ melanomas (n=3), and a significant increase in LTRs was found in Z18^trunc^-expressing tumors (P=4.5e-12, Wilcoxon signed-rank test) (Fig. 2A). Significant increases were also seen for LINE and SINE elements, although the magnitude of change was smaller (Table S2). To identify TEs that were most affected by Z18^trunc^ expression, we plotted the average expression difference by the average fold change (Fig. S2C) and the 27 elements with the largest expression difference were manually checked for autonomous expression using IGV (*29*). The most upregulated element was the ERV BHIKHARI-2, which is also known as crestin and is transiently expressed in the neural crest during development (*30*) and then re-expressed during melanoma initiation (*31*). Of the top upregulated TEs, 79% were LTRs (21/27), with 14 of these being Bhikhari-2 ERV elements. Of the TEs, 56% were expressed autonomously (15/27), i.e. independently of nearby genes, including all 14 BHIKHARI-2 elements and one other ERV (examples in Fig. 2B, S2D). To test whether other NEXT target RNAs were affected by Z18^trunc^ expression, we conducted ribo-minus RNA-seq of Z18^trunc^ (n=3) vs. eGFP (n=3) Triples melanomas (Fig. S2E). eRNAs and PROMPTs did not accumulate in Z18-expressing melanomas (Fig. S2F). These data show that Z18^trunc^-expression specifically promotes accumulation of ERV RNA, but not all NEXT targets (*3*), suggesting that Z18^trunc^ specifically regulates ERV messages.

**Figure 2.**
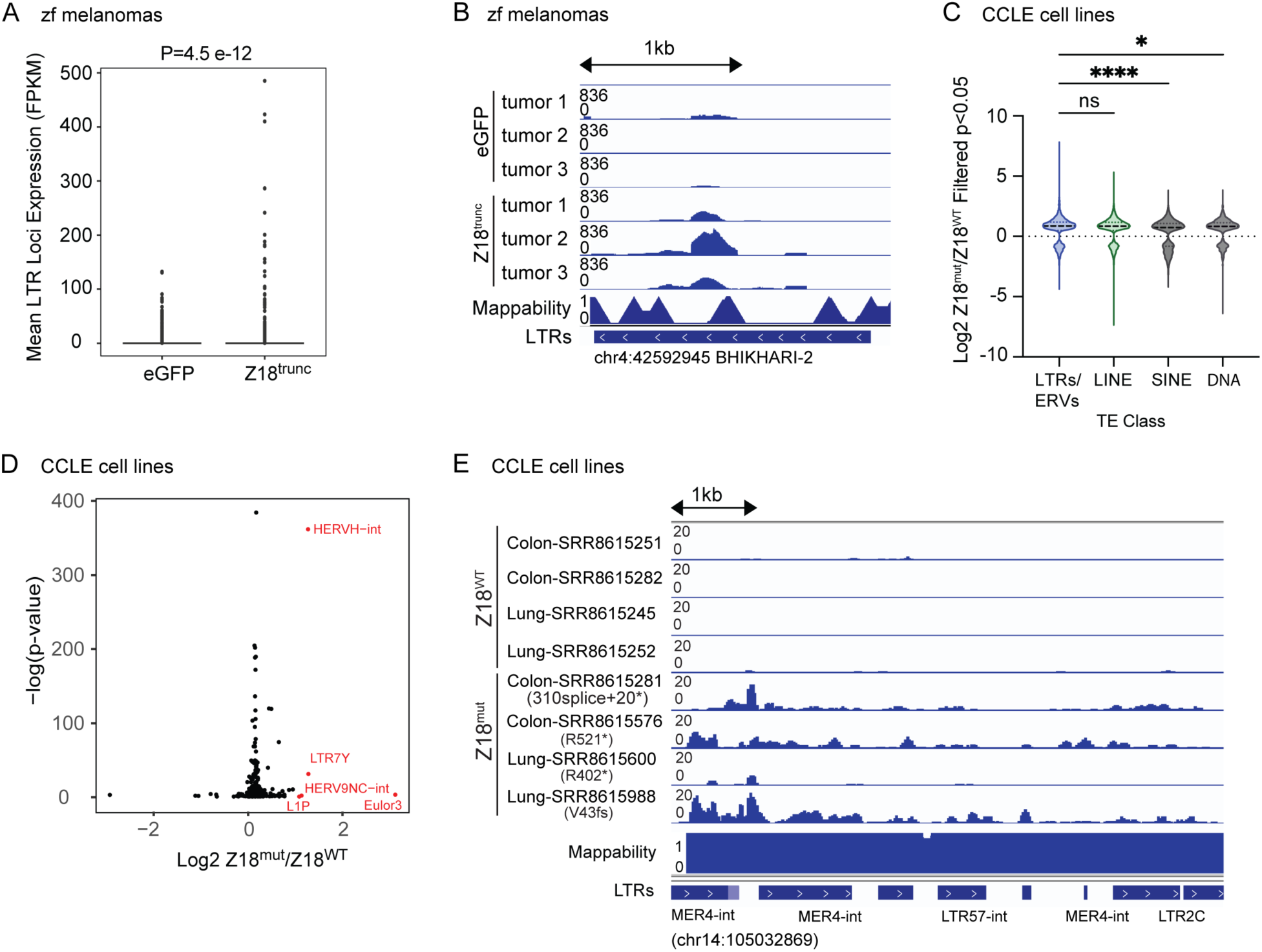
ZC3H18^trunc^ Promotes ERV Accumulation. A) Mean LTR loci expression (FPKM) from pA-selected RNA-seq from zebrafish melanomas expressing eGFP (n=3) vs. Z18^trunc^ (n=3). P=4.5e-12 (Wilcoxon signed rank test with continuity correction). B) IGV plot of an example LTR (ERV) in eGFP- and Z18^trunc^-expressing zebrafish melanomas. C) Log2 fold change of significantly differentially expressed (P<0.05) TEs in Z18^mut^ (n=11) vs. WT (n=11) CCLE cell lines. * P=0.018, **** P<0.001 (Ordinary one-way ANOVA). Thick dotted line = median. Light dotted line = quartiles. D) Volcano plot of expressed TE subclasses from Z18^mut^ (n=11) vs. WT (n=11) CCLE cell lines. Red labels = log2 fold change ≥1. E) IGV plot of significantly upregulated LTR (ERV) in Z18^mut^ CCLE cell lines.

As Z18 is evolutionarily conserved between zebrafish and humans, we wondered if Z18 might have a conserved role in regulating ERV accumulation. To test if Z18 plays a conserved role in regulating TE degradation, we utilized SQuIRE to quantify TE expression from publicly available pA-selected RNA-seq data. Cancer Cell Line Encyclopedia (CCLE) (*32*) cell lines with deleterious Z18 mutations (Z18^mut^) (n=11) were compared to cancer-type-matched cell lines with intact Z18 (n=11) (Fig. S2G, Table S3). Average log2 fold change in TE expression was plotted for TEs with a significant difference (P<0.05) between Z18^mut^ and control cell lines (Fig. 2C). All TE classes were found to be upregulated in Z18^mut^ lines as compared to controls, with LTRs being the most affected (Table S4). To determine which LTR subclasses were most affected, the average log2 Z18^mut^/Z18^WT^ subclass was plotted by the -log of the P-value (Fig. 2D) and average subclass expression difference between Z18^mut^ and control was plotted by the average log2 fold change (Fig. S2H). The most affected ERV subfamilies included HERVH-int, LTR7Y, and HERVK3-int. Two of the most significantly upregulated autonomously expressed ERVs, chr14:105032869 MER4-int (hereafter chr14 MER4-int) and chr13:109265101 HERVH-int, initiated in one element with transcription continued through several additional annotated element intervals before terminating (Fig. 2E, S2I). These data are consistent with a model wherein Z18 plays an important and evolutionarily conserved role in specifically suppressing ERV RNAs.

### ZC3H18^trunc^ Stabilizes ERV RNA in a Dominant Negative Manner

To directly test whether Z18^trunc^ mutations functionally promote ERV RNA accumulation, we built isogenic clonal Z18 mutant human melanoma cell lines that were heterozygous for Z18 patient truncating mutations (Z18^trunc/+^) (Fig. 3A, S3A-E, Table S5) (n=3) or Z18^WT^ (n=2). To introduce Z18^trunc/+^ mutation, CRISPR with homology-directed repair was pursued. We recovered one Z18^trunc/+^ clone with our homology-directed DNA mutation and two clones that harbored other Z18^trunc/+^ patient mutations. To determine if Z18^trunc^ exerts its effects by interfering with Z18^WT^, human melanoma cells with Z18 heterozygous loss-of-function mutations (Z18^-/+^) were also built (n=3) (Fig. S3F-G). We were unable to retrieve viable homozygous loss-of-function clones (0/23 clones with mutations), so transient bulk CRISPR was pursued (Fig. S3H-I) to assess the Z18 loss-of-function phenotype. Live adherent cells were collected for bulk Z18 CRISPR soon after protein depletion. The guide RNA used to make Z18^trunc/+^ clonal cell lines resulted in 83.3% of edited cell clones with out-of-frame truncating mutations (20/24) (Fig. 3B) while the two gRNAs used to make heterozygous loss-of-function clones (gRNA1 and 2) resulted in fewer out-of-frame mutations (10/23, 43.5%) (Fig. 3C) (gRNA locations in Fig S3A), suggesting that there may be selection for Z18^trunc/+^ mutations.

**Figure 3.**
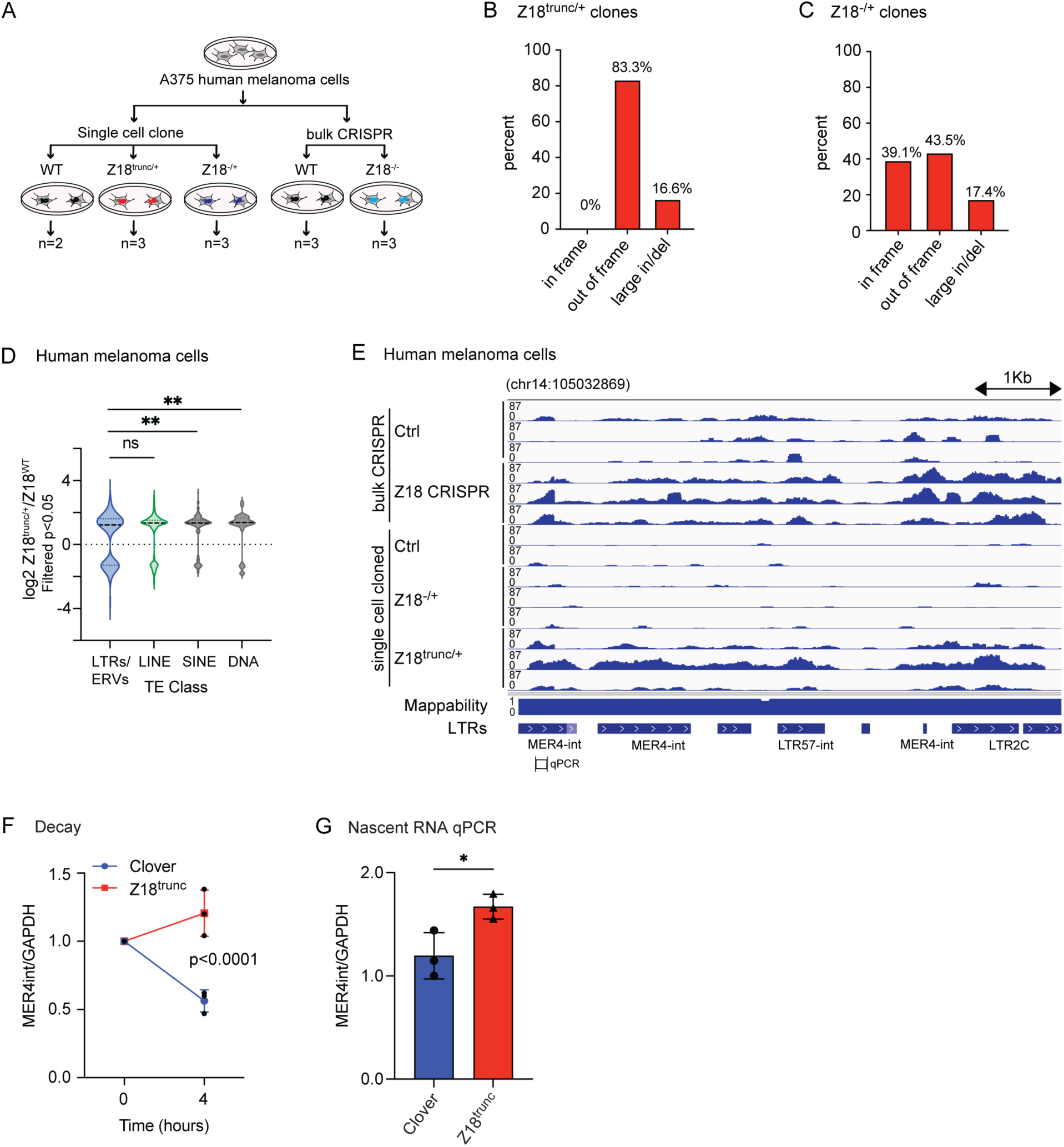
ZC3H18^trunc^ Stabilizes ERV RNA in a Dominant Negative Manner. A) Schematic of Z18 allelic series generated in A375 human melanoma cells. B-C) Percent of in-frame or out-of-frame Z18 CRISPR mutations for B) Z18^trunc^ clones (n=24) or C) Z18 loss-of-function clones (n=23). D) Log2 TE expression from Z18^trunc/+^ (n=3) vs. Z18^WT^ (n=2) human melanoma cell lines (filtered for P<0.05). LTR vs. SINE ** P=0.0024, LTR vs. DNA ** P=0.007, ns = non-significant (ordinary one-way ANOVA). Thick dotted line = median. Light dotted line = quartiles. E) IGV plot of significantly upregulated chr14 MER4-int LTR (ERV) in genetically modified cells lines from Fig. 3A. F) Decay of chr14 MER4-int using 4sU pulse-chase qPCR for Z18^trunc^- (n=3) vs. Clover-expressing (n=3) human melanoma cells (qPCR location in Fig. 3E). Mean ± SD. At 0 hours, non-significant. At four hours P<0.0001 (two-way ANOVA). G) qPCR of 4sU-labeled nascent chr14 MER4-int transcript expression (qPCR location in Fig. 3E) from Clover-and Z18^trunc^-expressing human melanoma cells. Mean ± SD. * P=0.032 (unpaired t-test, two-tailed).

To determine if Z18^trunc^ regulates ERV RNA in a dominant negative manner, SQuIRE was used to quantify TEs from pA-selected RNA-seq from the above Z18 allelic series. Average log2 fold change in TE expression was used to visualize significant changes (P<0.05) between Z18^trunc/+^ and Z18^WT^ cell lines. Consistent with our previous findings, LTRs and LINEs accumulated in Z18^trunc/+^ and bulk Z18 CRISPR-deleted cell lines as compared to the appropriate control cell lines (Fig. 3D, S3J). The most significantly upregulated autonomously expressed LTR in the Z18^trunc/+^ cell lines was the same chr14 MER4-int transcript that was regulated by Z18^mut^ in the CCLE cell lines (as in Fig. 2E; Fig. S3K, Table S6). Z18 loss-of-function via bulk CRISPR-deletion also resulted in chr14 MER4-int upregulation (Fig. 3E, top 6 rows), despite most transcripts being downregulated/degraded as cells undergo apoptosis(*33, 34*). Importantly, Z18^trunc/+^ enhanced chr14 MER4-int accumulation more than loss of one Z18 copy (Fig. 3E, last six rows), suggesting that Z18^trunc^ interferes with the ability of Z18^WT^ to clear ERV RNA, i.e. has dominant negative activity. Z18^trunc/+^ also similarly affected the next most significantly regulated and autonomously expressed LTR/ERV, chr16:35516910 HERVK3-int (hereafter chr16 HERVK3-int) (Fig. S3L, S3M). These data are consistent with Z18^trunc/+^ working in a dominant negative manner to promote accumulation of ERV RNA.

To determine how Z18^trunc^ mutations affect the broader transcriptome, Gene Set Enrichment Analysis (GSEA) (*35*) was performed on differentially expressed RefSeq transcripts from pA-selected bulk RNA-seq of the Z18^trunc/+^ vs. control single-cell clones. Of the Hallmark signatures(*36*), the most significantly upregulated pathway was “TNFA_SIGNALING_VIA_NFKB” (Fig. S3N, FWER P=0, NES 2.44) (Fig. S3N, Data S2, first tab) and the two most upregulated pathways from the chemical and genetic perturbations datasets were associated with viral infection, suggesting viral mimicry (“RESPIRATORY_SYNCYTIAL_VIRUS_INFECTION_A594_CELLS_UP” and “HUMAN_PARAINFLUENZA_VIRUS_3_INFECTION_A594_CELLS_UP”) (Fig. S3O, Data S2, second tab). We hypothesize that widespread ERV RNA accumulation promoted increased expression of anti-viral genes, as has been reported for transcriptional derepression of ERVs (*17, 37, 38*).

To determine if the increase in the chr14 MER4-int transcript was due to increased transcription or defective degradation, we measured its transcription and decay kinetics. Due to the heterogeneity produced by CRISPR non-specific targeting and single-cell cloning and because Z18^trunc^ functions in a dominant negative manner, Z18^trunc^- and Clover-expressing human melanoma cell lines were generated (Fig. S3P). We confirmed upregulation of the chr14 MER4- int transcript in the Z18^trunc^-expressing human melanoma cell lines using qPCR (Fig. S3Q, qPCR location in Fig. 3E), while levels of chr16 HERVK3-int were unchanged (Fig. S3R, qPCR location in Fig. S3M). We measured the decay of chr14 MER4-int in Clover- and Z18^trunc^-expressing cell lines using 4sU pulse-chase and found significant stabilization of chr14 MER4-int in Z18^trunc^- expressing (n=3) as compared to Clover-expressing (n=3) cell lines (Fig. 3F). To assess chr14 MER4-int transcription, nascent RNA(*39*) qPCR was completed for Clover- and Z18^trunc^-expressing human melanoma cells, which showed that chr14 MER4-int was also being transcribed at higher levels (Fig. 3G), consistent with a report that LINE1 chromatin is more accessible upon depletion of ZCCHC8, a core NEXT component (*9*). This data suggests chr14 MER4-int decay could play a role in its own chromatin silencing. Together our data are consistent with the model that Z18^trunc^ acts to stabilize ERV messages in a dominant negative manner.

### ZC3H18^trunc^ Directly Promotes Oncogenic ERV RNA Stabilization

To determine the mechanism of Z18^trunc^-mediated oncogenesis, we identified the interacting partners of full length (Z18^fl^) and truncated Z18 via immunoprecipitation mass spectroscopy (IP-MS). V5-tagged Z18^fl^ and Z18^trunc^ were transiently expressed in A375 human melanoma cells. Nuclear protein was extracted under native conditions and Z18 was IPed in triplicate using a V5 antibody. IPs were done in the presence or absence of RNase A/T1 to define RNA-dependent and -independent protein interactions (Fig. 4A). Data were filtered to include proteins that were >4.5× enriched over a published control IP (*6*) and had at least 5 total peptides measured in each replicate in the most permissive condition (Data S3). Z18 abundantly bound NEXT components, ZCCHC8 and MTR4, consistent with our data showing that Z18 regulates ERV RNA turnover, which requires NEXT (*10*).

**Figure 4.**
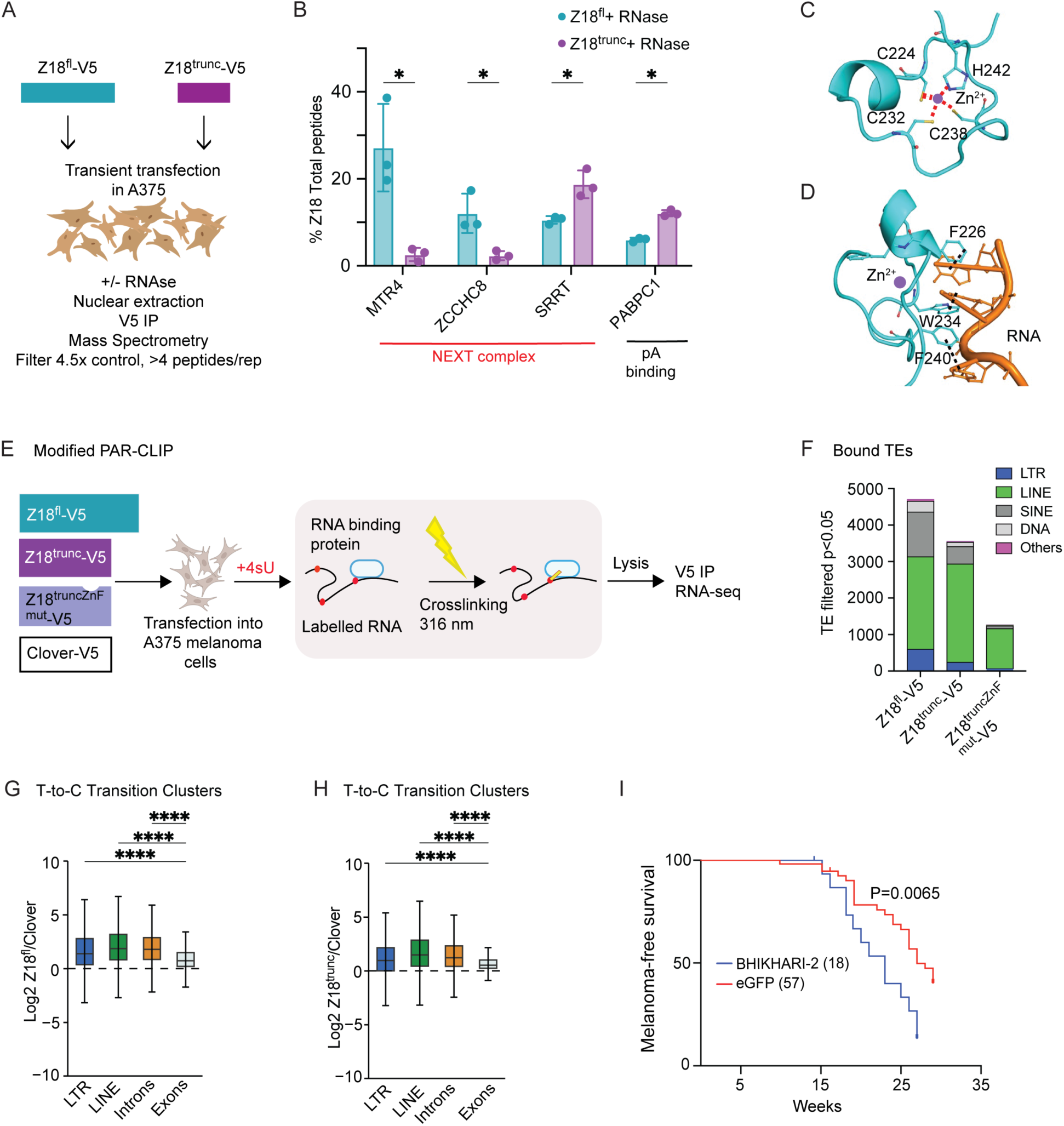
ZC3H18^trunc^ Directly Promotes Oncogenic ERV RNA Stabilization. A) Schematic of immunoprecipitation-mass spectrometry (IP-MS). fl = full length. B) IP-MS total peptides normalized to Z18 total peptides for proteins with differential binding between Z18^fl^ vs. Z18^trunc^ (+RNase). MTR4 * P=0.028, ZCCHC8 * P=0.043, SRRT * P=0.046, PABPC1 * P=0.010; (two-sided t-test). C-D) Modeled structure of the Z18 zinc finger domain (cyan) with C) Zn^2+^ ion (purple) coordinated with three cysteines and a histidine and D) RNA (orange) with dashed lines indicating predicted pi-pi stacking interactions. E) Schematic of photoactivatable ribonucleoside-enhanced crosslinking and immunoprecipitation (PAR-CLIP) workflow. F) Significantly (P<0.05) bound TEs from Z18^fl-^V5, Z18^trunc^-V5, and Z18^truncZnFmut^-V5 vs. Clover-V5 control from SQuIRE PAR-CLIP analysis. G-H) Log2 Z18^fl^ or Z18^trunc^ vs. Clover binding to LTR-, LINE-, intron-, or exon-containing clusters. In box plots, the black horizontal line indicates the median, the box covers the interquartile range (IQR), and the whiskers extend to 1.5× the IQR. Medians and two-sided Wilcoxon rank sum P-values in Table S7. I) Percent melanoma-free survival for Triples zebrafish with melanocyte-specific expression of eGFP or the ERV BHIKHARI-2. P=0.0065 (log rank). n= zebrafish.

To determine how Z18^trunc^ stabilizes ERV RNA, we evaluated Z18^fl^ and Z18^trunc^ binding partner differences (Fig. S4A). As expected, the loss of the C-terminal RNA surveillance recruitment domain resulted in a significant loss of MTR4 and ZCCHC8 binding to Z18^trunc^ as compared to Z18^fl^ (Fig. 4B, S4B). Four proteins with homology to splicing factors also had significantly reduced binding to Z18^trunc^ (FNBP4, PRPF4B, RBM25, and PRPF40A) (Fig. S4C). In contrast, PABPC1 and SRRT were observed with stable to increased binding to Z18^trunc^ (Fig. 4B, S4D). PABPC1 is a polyadenosine-binding protein that tends to be amplified in cancer (*40*) and has been shown to shield RNAs from decay(*41*). PABPC1 is essential for LINE1 retrotransposition (*42*) but has not been reported to have a function in NEXT and SRRT is a cap-binding protein associated with the NEXT complex. These data indicate that Z18^fl^ interacts with the NEXT complex and that Z18^trunc^ has reduced binding to NEXT while maintaining interactions with two RNA-binding proteins, PABPC1 and SRRT. These findings nominate a mechanism for how Z18^trunc^ promotes ERV RNA stabilization and are consistent with reports of the loss of NEXT components causing ERV upregulation (*9, 10*).

To consider how Z18^trunc^ exerts its dominant negative activity, we proposed two models: Z18^trunc^ could bind and sequester 1) Z18^WT^ from its RNA substrates (Fig S4E, Model 1) or 2) substrate RNAs away from Z18^WT^ (Fig. S4E, Model 2). In Model 1, Z18^trunc^ should be predominantly localized in the same compartment as Z18^fl^ and should efficiently bind Z18^fl^. We previously noticed by IP-MS that Z18^trunc^ was found at lower levels in the nuclear fraction than Z18^fl^ (Fig. S4F). We hypothesized that Z18^trunc^ was being exported to the cytoplasm, as was previously seen for a C-terminal mutant of Z18 (*23*). Immunoblotting for V5-tagged Z18 indicated that Z18^fl^ was mostly found in the nucleus as expected, while Z18^trunc^ was predominantly found in the cytoplasm (Fig. S4G) where it continued to associate with SRRT and PABPC1 (Fig. S4H). As Z18^trunc^ is predominantly cytoplasmic, it is unlikely that sequestration of nuclear Z18^fl^ explains its dominant negative function.

To further test whether Z18^trunc^ could sequester Z18^WT^ from its substrate, we asked whether Z18^trunc^ bound Z18^fl^. Z18 IP-MS unique peptide measurements were plotted along the protein length (Fig. S4I). As expected, Z18^trunc^ samples (+/- RNase) had only a few peptides measured distal to the truncation site, suggesting that Z18^trunc^ does not bind (or inefficiently binds) Z18^fl^. To confirm that Z18^trunc^ does not sequester Z18^fl^, we performed IP-westerns of Z18^trunc^ from the nuclear and cytoplasmic fraction for presence of full-length Z18 and found no signal (Fig. S4J-K). These data show that Z18^trunc^ is not enriched in the correct compartment and does not have detectable Z18^WT^ binding, making Model 1 unlikely.

We next considered whether Z18^trunc^ might exert its dominant negative activity by binding and sequestering ERV RNA from the NEXT complex (Fig. S4E, Model 2). Since the ability of Z18 to regulate retroelement RNA is highly evolutionarily conserved, we analyzed Z18’s homology to determine how Z18 could bind RNA. We found that Z18 has two highly conserved domains, including the nuclear RNA surveillance binding domain and a highly conserved zinc-finger domain (Fig. S4L). While zinc-finger domains are ubiquitous in the eukaryotic proteome, Z18 is one of 57 human proteins harboring the C-x-C-x-C-x-H (CCCH hereafter) motif (*43*). CCCH proteins directly bind and regulate RNA, and many have been implicated in immunologic pathways (*43*). We used the RBM22 (*44*) (PDB 6ID1) structure to create a homology model of the Z18 zinc-finger domain (Fig. S4M) as the Z18 structure is not known. Our model predicted that the CCCH zinc-finger domain has a Zn^2+^ ion coordinating with C224, C232, C238, and H242 (Fig. 4C), and that the highly conserved aromatic residues F226, W234, and F240 directly bind RNA via pi-pi stacking interactions (Fig. 4D). We hypothesized that Z18^trunc^ binds RNA via the aromatic amino acids in the zinc-finger domain.

To determine whether Z18 directly binds ERV RNA via the aromatic amino acids in the zinc-finger domain, we used photoactivatable ribonucleoside-enhanced crosslinking and immunoprecipitation (PAR-CLIP) (*45*) to determine the RNAs that bind Z18. V5-tagged Clover, Z18^fl^, Z18^trunc^, and Z18^trunc^ ZnF aromatic mutant (hereafter Z18^truncZnFmut^) were transiently expressed in A375 human melanoma cells (n=3 each) and IPed. Bound RNA was sequenced and Z18 differential binding to TEs as compared to Clover was measured using SQuIRE (Fig. 4E, S4N-P). Z18^fl^ significantly bound many TEs (4702), Z18^trunc^ maintained binding to most TEs (4134), while Z18^truncZnFmut^ bound fewer TEs (1266) (P<0.05 from DESeq2) (Fig. 4F-G, S4Q-R). This finding shows that the aromatic residues in Z18’s zinc-finger domain are required to stabilize TE binding. Z18^trunc^ bound significantly more LINE elements than Z18^fl^, although RNA-seq showed that LTRs/ERVs were more highly regulated (Fig. 3D). As species-specific LINE-1 elements can mobilize in cancer, we investigated whether LINE-1 retrotransposition could be contributing to Z18^trunc^ cancer phenotypes. Using catalogs of published somatically-acquired LINE-1 insertions in a pan-cancer study (*46*), we found that there was no statistical increase in LINE-1 retrotranspositions in Z18-mutated cancers (average Z18^trunc^ 10.09, n=11; average rest 6.57, n=326; P=0.72, two-sided t-test). We also found no increase in LINE-1 open reading frame 1 (ORF1) protein expression in the Z18^trunc^-expressing cell lines (Fig. S4S). These data shows that Z18^fl^ and Z18^trunc^ can directly bind LINEs and LTR/ERV messages and that the aromatic residues in the Z18 zinc-finger domain are required to stabilize TE binding.

To determine the specific RNA sequences bound by Z18 in the PAR-CLIP data, we utilized WavCluster (*47*), which identifies high confidence binding events from clusters of T-to-C transitions (hereafter clusters). Z18^fl^ and Z18^trunc^ had a similar number of clusters (296229 and 284329, respectively) while Clover resulted in fewer clusters (144336), which represent background RNA binding. To determine the identity of Z18-bound RNAs, clusters were annotated for LTRs, LINEs, introns, and exons (Fig. S4T-W). We found that Z18^fl^ and Z18^trunc^ clusters overlapped with more LTRs, LINEs, and introns as compared to exons. Bound RNA (from Z18^fl^ cluster regions) was quantified for Z18^fl^ and Z18^trunc^ vs. Clover PAR-CLIP (Fig. 4G-H, statistics in Table S7). This quantification showed that Z18^fl^ and Z18^trunc^ had significantly enriched binding to LTRs, LINEs, and introns as compared to exons (example Z18-bound transcript with intron inclusion in Fig. S4X). As introns (*48*) and retroelement (*49*) messages have higher AU nucleotide content than exons, we next investigated whether Z18 bound AU-rich RNA as compared to background Clover-bound RNA. We found that Z18^fl^ and Z18^trunc^ bound more AU-rich RNA than Clover. This effect was decreased for the Z18 aromatic zinc-finger mutant (Fig. S4Y). These analyses show that Z18^fl^ and Z18^trunc^ directly bind and regulate retrotransposon RNA via conserved aromatic amino acids in the zinc finger domain, supporting that Z18 exerts its dominant negative activity via binding RNA.

We wondered whether ERV RNA that is stabilized by Z18^trunc^ mutations, could directly contribute to oncogenesis. We selected one of the most highly expressed ERVs from Z18^trunc^-expressing zebrafish melanomas, chr4:42592945 BHIKHARI-2 (hereafter BHIKHARI-2) (Fig. 2B) and expressed this element in the zebrafish system. Notably, expression of BHIKHARI-2 was sufficient to expedite melanoma onset in the zebrafish system (Fig. 4I, S4Z). This BHIKHARI-2 ERV is a solo LTR, i.e. it underwent LTR recombination, resulting in loss of intervening protein coding regions. Thus, we predict that BHIKHARI-2 RNA directly contributes to oncogenesis not via a resulting protein product as this message does not contain known coding sequence. This data shows that ERV RNA itself can functionally contribute to oncogenesis.

## Discussion

By studying the effects of Z18 mutations, we have revealed a functional role for ERV RNA expression in melanoma. Our data suggest that Z18 mutations are selected for in melanoma and that Z18^trunc^ mutations inhibit Z18^WT^’s evolutionarily conserved role of regulating ERV RNA turnover. As Z18 truncating mutations promote higher ERV RNA accumulation compared to loss of one Z18 copy, our data are consistent with Z18^trunc^ interfering with the ability of Z18^WT^ to target ERV RNA for degradation. The cancer-associated Z18^trunc^ protein loses association with nuclear RNA surveillance machinery, is mislocated to the cytoplasm, and binds ERV RNA, protecting these messages from degradation. We found that ERV RNA itself was sufficient to expedite oncogenesis. ERV RNA has been long known to be highly expressed in cancer, but the causes and consequences have been elusive. Our data suggest that specific cancer-associated mutations can promote the accumulation of ERV RNA and that ERV RNA can directly contribute to oncogenesis.

We previously found that defective nuclear RNA surveillance of ptRNAs from protein-coding genes is oncogenic (*6*). Since we (*6*) and others (*4*) have identified Z18 as a component of the PAXT complex that degrades ptRNAs, we hypothesize that Z18^trunc^ mutations would result in accumulation of ptRNAs. Instead, we found that truncating Z18 mutations resulted in a specific accumulation of retroelement RNA in zebrafish, CCLE cell lines, and engineered human melanoma cells, with ERVs being most affected. As Z18 is also a component of NEXT (*10*) which normally degrades ERVs, Z18’s specificity in regulating ERV turnover could help to reveal how NEXT targets substrate RNAs for degradation. This work suggests that disruption of Z18-mediated RNA surveillance is similar to the CDK13-mediated pathway (*6*) as there is accumulation of polyadenylated RNA, but the identity of the accumulated RNA is distinct. As both mutations are oncogenic, this work further supports the idea that loss of RNA quality control is a hallmark of cancer cells.

Although Z18’s role in cancer has not been investigated previously, there is one report of a patient with severe congenital neutropenia that transformed into acute myeloid leukemia. This patient harbored a Z18^trunc^ mutation (777fs) in the offending clone (*50*), supporting that although these mutations are rare, they are likely functional. We found that Z18 mutations are statistically enriched just upstream of the domain required to recruit nuclear RNA degradation machinery, suggesting a role beyond loss-of-function. We found Z18^trunc/+^ enhanced ERV RNA accumulation more than loss of one Z18 copy, demonstrating dominant negative activity. We previously found that CDK13 mutations also work in a dominant negative manner (*6*), which may represent a common mechanism for mutations related to nuclear RNA surveillance. We hypothesize that dominant negative mutations are sufficiently severe to perturb RNA metabolism while avoiding lethality induced by a full loss-of-function.

Sorting of nuclear transcripts is an essential cellular process, with capped, spliced, and polyadenylated transcripts being exported and aberrant or unmodified RNAs being degraded in the nucleus. RNA fate in the nucleus is often binary: either RNAs are identified by export machinery and are stabilized, or they are actively targeted for degradation. SRRT sits at the nexus of nuclear RNA fate (*51, 52*), as messages bound to the SRRT/Z18 complex are degraded by the nuclear RNA exosome, while SRRT bound with the export protein PHAX stabilizes and exports transcripts (*53*). We propose that Z18^trunc^-containing complexes escape to the cytoplasm, protecting bound ERV RNA from degradation.

Z18, which is named for its CCCH zinc-finger domain (<u>ZC3H</u>18), is one of nearly 60 human proteins containing a CCCH zinc-finger (*43*). CCCH zinc-fingers have been shown to bind to RNA, and many, including Z18, have functional roles in immune responses (*43, 54*). Indeed, we found that Z18 normally specifically bind ERV RNA and targets it for degradation. As retroelements comprise almost half of the human genome (*11, 55*), perhaps it is not surprising that retroelement transcription can affect cellular phenotypes, even when immobile (*12*). Across evolution, multiple organisms have evolved and retained effective mechanisms to suppress ERV expression, even for immobile elements. The most well-described method of suppressing ERV expression is direct silencing executed by Krab zinc finger proteins (KZFPs) and other DNA-binding zinc finger proteins. The role of KZFPs in limiting transposable element expression is so profound in evolution and multiple studies have shown that TE evolution drives host zinc finger evolution (*56–61*). Our work highlights that RNA-binding zinc fingers may play a complementary role to DNA-binding zinc fingers by activating degradation of ERV RNAs that escape transcriptional repression.

The human silencing hub (HUSH) complex has also been shown to repress expression of intron-less retroelements (*62*). Although ZCCHC8, a component of the NEXT complex, directly interacts with HUSH (*10*), Z18 was not detected with any of the key HUSH complex members in prior work (*23*) or in our IP-MS experiments (except for PPHLN, which was below our filtering thresholds). Instead, we saw enrichment of proteins related to splicing and found that Z18 tends to bind intron-containing messages. For example, we observed that two of the most regulated ERV-containing messages chr14 MER4-int and chr16 HERVK3-int underwent splicing with either another retroelement or downstream host sequences. Spliced ERV RNAs, which are not regulated by HUSH silencing, may require NEXT targeting via Z18, which could explain Z18’s specificity for TE targets over unspliced RNAs such as PROMPTs and eRNAs.

Retroelements, especially LTR/ERVs (*16, 17*), have been long known to be expressed at increased levels in multiple cancer types, suggesting that the production of RNA from TE loci is under positive selection. The upstream causes and downstream effects have remained elusive. Intriguingly, studies have found that proper ERV message levels are required for early embryonic development with both upregulation (*63*) and downregulation (*64*) causing embryonic lethality. As many molecular processes required in development are hijacked by cancer, this literature suggests that despite their immobility, ERV RNA expression could be repurposed in human cancer cells to promote oncogenesis. Indeed, we found expression of a Z18-target solo ERV was sufficient to expedite melanoma onset in a zebrafish model, showing that aberrant ERV expression itself can contribute to oncogenesis. More studies are needed to identify the mechanism by which ERV RNA expression directly contributes to oncogenesis.

Z18 mutations via ERV RNA accumulation may also have important therapeutic implications for patients. We observed that Z18 mutations activated anti-viral gene expression, similar to what has been observed for ERV transcriptional activation (*17, 37, 38*) which has been shown to result in immune infiltration, increased antigenicity, and potentiation of immune therapy responses in multiple cancers (*17–19*). In one notable manuscript, ERV-encoded antigens were shown to be a predictive marker for immune therapy responses in lung cancer patients (*20*). We hypothesize that patients with Z18-mutant cancers may be more likely to respond to immune therapy due to ERV message stabilization, either through innate immune activation, RNA dependent stress, or translation of ERV-encoded peptides. In addition, multiple studies have revealed thousands of cancer-specific transcripts that result from noncanonical splicing between LTRs/ERVs and host exons (*65, 66*) with many pan-cancer antigens (*15*). As host exon to TE spliced transcripts are the RNA type that we observe to be most regulated by Z18, there could be a role for temporary Z18 inhibition in Z18^WT^ cancers in order to elicit immune therapy responses. Although Z18 mutations themselves are rare, there are therapeutic implications for this pathway.

We found that most Z18 mutations occur in microsatellite stable tumors, although Z18^trunc^ and Z18 non-truncating were enriched in microsatellite unstable tumors as expected. Microsatellite unstable tumors are known to be more responsive to immune therapy (*67–69*), however, not all patients benefit. In one colon adenocarcinoma study, high ERV expression was found to be an independent predictor from MSI-status of immune activation and CD8+ T cell infiltration (*70*). Mutated Z18 could therefore be a predictive marker for immune therapy responses in cancers with mutated or loss of Z18, regardless of MSI-status.

We propose that Z18 plays an evolutionarily conserved role in identifying and destroying ERV messages that escape transcriptional silencing, in contrast, Z18^trunc^ mutations promote specific retroelement RNA stabilization. This is the first example to our knowledge of a genetic mechanism contributing to ERV accumulation in cancer. This is also the first example of which we are aware that shows that ERV RNA expression itself can contribute to oncogenesis. Our work further supports the concept that lack of proper nuclear RNA degradation shapes the cancer transcriptome, contributing to oncogenesis as well as nominating therapeutic vulnerabilities.

## Acknowledgments

The results shown here are in part based upon data generated by the TCGA Research Network: https://www.cancer.gov/tcga. We would also like to acknowledge the following funding sources: National Cancer Institute K08 CA248727 (MLI) National Cancer Center Postdoctoral Award (TT) Ladieu Family Melanoma Research Fund (MLI) King Family Fund for Melanoma Research (MLI)

## Author contributions

Conceptualization: TT, AMC, KHB, PLB, MLI

Methodology: TT, AMC, PS, XH, PLB, MLI

Formal Analysis: JK, PS, XH, SB, JB, TZ, PLB, MLI

Investigation: TT, AMC, MLI

Writing – original draft: TT, AMC, MLI

Writing – review & editing: TT, AMC, MLI

Supervision: KMB, SJC, DL, KHB, PLB, MLI

Funding acquisition: TT, KHB, PLB, MLI

## Competing interests

Authors declare that they have no competing interests.

## Data and materials availability

The datasets generated and/or analyzed during the present study have been uploaded to the Gene Expression Omnibus (GEO pending). The CCLE data are publicly available. All zebrafish strains, human cell lines, and DNA vectors are readily available through the corresponding author. Where possible, we can share the remaining zebrafish tumor material; however, this material is limited in abundance. All antibodies are commercially available.

## Supplementary Materials

Materials and Methods

Figs. S1 to S4

Tables S1 to S7

References 1-32

Data S1 to S3

## Notes

### Competing Interest Statement

The authors have declared no competing interest.

