## Supplemental file for "Recurrent oncogenic ZC3H18 mutations stabilize endogenous retroviral RNA"

**The file includes:**

Materials and Methods
Figs. S1 to S4
Tables S1 to S7
References #1-32

**Other Supplementary Materials for this manuscript include the following:**

Data S1 to S3

#### Materials and Methods

##### Patient Data Analysis:

A dataset of non-redundant studies spanning various cancer types from cBioPortal was compiled. In total, 85 unique studies were identified (Data S1, "Pan-cancer Patient Sample Refs"), encompassing 23411 samples, each containing at least one sample with mutations in the *ZC3H18* (*Z18*) gene. Duplicate or serial samples from the same individual were excluded. Amino acid residues with >5 mutations were plotted. To analyze whether truncating mutations are enriched upstream of the RNA surveillance recruitment domain, a Fisher's exact test was used (Data S1 "$Z18^{trunc}$ Fisher Exact Test"). MSIsensor[1] scores were used to identify microsatellite unstable samples within the TCGA PanCancer dataset[2]. Samples with Z18 truncating and non-truncating mutations were compared to Z18 wild-type ($Z18^{WT}$) samples to determine which were associated with microsatellite instability (MSIsensor >10) using an odds ratio and a Fisher's exact test (two-sided) (Table S1).

##### Zebrafish Experiments:

**Cloning:** The MiniCoopR (MCR) Gateway system was used[3]. MCR overexpression constructs were assembled using Gateway vectors including the *mitfa* promoter, relevant coding sequence, 3′ poly-A (pA) tail, and *MCR* destination vector with LR Clonase II Plus (Thermofisher, 12538120). The human Z18 coding sequence was obtained from Dharmacon (Accession: MHS6278-202759301 Clone ID: 6042036). Truncated Z18 was cloned with the R680Gfs*5 (hereafter $Z18^{trunc}$) patient mutation using *in vitro* mutagenesis (NEB, E0554S). A gene fragment for the chr4:42592945 BHIKHARI-2 ERV (hereafter BHIKHARI-2) was ordered from BioTwist and this fragment was assembled with SalI and EcoRI linearized empty MCR vector (NEB, E2621).

**Melanoma model:** Animal studies were approved by the Animal Care and Use Committee at the Dana-Farber Cancer Institute (Protocol 21-027). Experiments were performed as published[3,4]. ARRIVE guidelines were followed where possible[5]. Briefly, *$p53^{-/-}$; mitfa:$BRAF^{V600E}$; $mitfa^{-/-}$* (hereafter referred to as Triples) one-cell embryos (randomized) were injected with either 20 ng/µL control or experimental DNA along with *tol2 in vitro* transcribed RNA for integration. In all experiments, DNA which overexpressed the gene of interest was marked with a *mitfa* mini gene that rescues Mitfa, allowing cell autonomous melanocyte genetics (schematic in Fig. S1A). Control vectors expressed eGFP. Embryos were sorted for melanocyte rescue (vector integration) at 5 days post fertilization. We aimed to generate 20 fish per genotype to give us a 75% chance of detecting changes in tumor onset (hazard ratio of .30, alpha 5%). The same number of zebrafish were raised in control and experimental tanks to control for density effects. Zebrafish with >10% of their body length melanocyte rescue at 9 weeks were scored for the emergence of raised melanoma lesions greater than or equal to 1 $mm^2$. Melanoma-free survival curves and log-rank tests were generated in Prism. Black patches that disrupted normal stripes and had a diameter greater than the distance between that zebrafish's eyes were quantified at 9 weeks.

**Western blotting:** Protein was quantified using the DC Protein Assay (Biorad, 5000116). The following antibodies were used: anti-Z18 antibody (Novus, NBP1-82570, 1:1000), anti-beta actin (CST, 3700S, 1:1000), anti-GFP (Santa Cruz, sc-9996, 1:1000). Rabbit or mouse secondary HRP antibodies were incubated for 1 hour at room temperature (20-25°C) (CST, 7074S or 7076S, 1:2000). Films were developed with Pierce ECL substrate (Thermofisher, 32106).

**RNA-seq:** Triples zebrafish melanomas from different individual fish with expression of eGFP (n=3) or $Z18^{trunc}$ (n=3) were collected. Tumors were homogenized, RNA was purified (Zymo Research, R1054), DNA was removed, and RNA was polyA-selected (NEB, E7490S). Additional eGFP (n=3) or $Z18^{trunc}$ (n=3) melanomas were processed similarly and ribo-depleted (NEB, E7405). All samples underwent stranded library preparation (NEB, E7760S) and were sequenced on a HiSeq 4000.

**RNA-seq analysis:**

*Custom genome mapping for transgene verification*: The raw sequence reads from pA-selected and ribo-minus RNA seq were trimmed using Cutadapt (v. 4.6) with the parameters: cutadapt -j 4 -q 30 -a GATCGGAAGAGCACACGTCTGAACTCCAGTCAC -a CTGTCTCTTATACACATCT -g AGATGTGTATAAGAGACAG -A GATCGGAAGAGCGTCGTGTAGGGAAAGAGTGTAGATCTCGGTGGTCGCCGTATCATT -A CTGTCTCTTATACACATCT -G AGATGTGTATAAGAGACAG -m 50 -e 0.02 [6]. The trimmed reads were mapped to a custom reference genome containing primary chromosomes of the *Danio rerio* genome build danRer11 (i.e., chr1-chr25) plus the human gene *Z18*. Mapping was performed using STAR (v. 2.5.3a) in paired-end mode with parameters --outSAMtype BAM SortedByCoordinate --quantMode GeneCounts --outWigStrand Stranded --outWigType bedGraph[7]. Bedgraph files were converted to bigwig format using bedGraphToBigWig[8]. Bigwig files were visualized in Integrative Genomics Viewer (IGV)[9].

*RNA-seq analysis for transposable elements (TE)*: TEs were measured from polyA-selected RNA-seq. SQuIRE[10] was used for mapping to the danRer11 genome, counting, and examining differential expression of individual TE as annotated by the RepeatMasker gtf (downloaded from UCSC on 9/20/2022). The average expression of each individual TE was calculated for all samples. Only loci that had expression on the correct strand in at least one sample were considered. In cases where an individual locus was expressed in a subset of samples, "NaN" values were updated to "0"s. A Wilcoxon signed rank test with continuity correction was used to determine statistical differences in TE expression between $Z18^{trunc}$ vs. eGFP conditions. Mean LTR loci expression (FPKM) were visualized in Fig. 2A. Statistics for other TE classes are included in Table S2. The average FPKM log2 fold change was plotted by the difference for all loci in Fig. S2C. Uniquely mapping reads for the most highly expressed TEs with a significant change were visualized in IGV[11].

*RNA-seq analysis of canonical NEXT targets:* Ribo-minus RNA-seq reads from eGFP or $Z18^{trunc}$-expressing zebrafish melanomas were mapped to *Danio rerio* dr11 genome build using STAR 2.5.3a[7] with the command parameters: STAR –runMode alignReads –twopassMode Basic –outFilterMutimapNmax 20 –outFilterMismatchNmax 999 –outFIlterMismatchNoverLmax 0.033 –alignIntronMin 70 –alignIntronMax 500000 –alignMatesGapMax 500000 –alignSJoverhangMin 8 –alignSJDBoverhangMin 1 –outSAMstrandField intronMotif –outFilterType BySJout. Bed files containing the genomic coordinates of known NEXT target RNAs were generated as follows. Enhancer regions were identified using a published H3K27ac ChIPseq bed file from a zebrafish melanoma cell line (GEO accession GSM1953851_CKK2_11_peaks.narrowPeak.gz)[4]. As this was a narrow peak file mapped to Zv9, liftOver from UCSC[12] was used to convert Zv9 coordinates to dr11. Using intersectBed –v function from Bedtools version 2.30.0[13], H3K27ac-enriched regions that excluded gene bodies were identified as intergenic enhancers, while H3K27ac-enriched regions that overlapped the gene body excluding the 200 nucleotides downstream of the transcription site (to eliminate promoters) were identified as intragenic enhancers. PROMPTs were identified as the 200 nucleotides upstream in the antisense direction of all annotated gene transcription start sites. The mapped reads falling within each region of the NEXT target bed files were counted using the featureCounts function of subread version 2.0.3[14]. Antisense strand specificity was required for PROMPTs and intragenic enhancers. Counts were input into DEseq2[15] for eGFP- vs. $Z18^{trunc}$-expressing samples.

#### Human Cell Line Experiments:

**CCLE RNA-seq analysis:** Eleven cell lines from the Cancer Cell Line Encyclopedia (CCLE)[16] with Z18 mutations (Table S3) and available RNA-seq (GEO PRJNA523380) were downloaded. To serve as controls, 11 cancer-type-matched CCLE lines that were $Z18^{WT}$ were selected (Table S3). SQuIRE[10] was run on fastq files. TE loci that were expressed in at least one sample were included in downstream analysis. "NA" values were then replaced with 0s. Significantly changed TEs were filtered for p-value <0.05 (paired Wilcoxon test) (n=49708). Average difference in TE subfamily expression was calculated (average $Z18^{mut}$ - average $Z18^{WT}$). Log2 fold change was calculated with a pseudocount of 0.00001. The -log p-value versus the log2 fold change (mutant / wild type) for each subfamily was plotted and the subfamilies with a log2 fold change greater than or equal to 1 were labeled (Fig. 2D). The average difference versus the log2 fold change for each subfamily was plotted and subfamilies with an FPKM increase >0.5 and a log2 fold change >1.75 were labeled (Fig. S2H).

**Human cell lines:** A375 human melanoma cells (ATCC CRL-1619) were identity-verified via Short Tandem Repeat analysis then used for transient transfections or stable line generation. Routine monthly mycoplasma testing was done with a PCR-based detection kit (Sigma, MP0025). All cell lines were mycoplasma negative. Cells were grown in DMEM with 4.5g/L D-glucose and L-Glutamine supplemented with 10% FBS and penicillin/streptomycin or selection antibiotics.

*Z18 heterozygous loss-of-function ($Z18^{-/+}$) and Z18 heterozygous patient-truncating ($Z18^{trunc/+}$) lines:* gRNAs were selected via CHOPCHOP14[17] and cloned into Gecko2.0 vectors (Addgene 49535) which express both Cas9 and the relevant gRNA in one plasmid. gRNA efficiency was assessed via bulk transient transfection (Invitrogen, 100022050) into A375 cells, 72-hour puromycin selection, and Z18 immunoblot. The most efficient gRNAs (>50% protein reduction) were selected for single cell cloning using limited dilution. To introduce $Z18^{trunc/+}$

mutation, CRISPR with homology-directed repair was pursued using co-transfection of a vector expressing an sgRNA targeted within 20 base pairs of the 680-mutation site and a single-stranded HDR donor DNA template (Integrated DNA Technologies). Three $Z18^{-/+}$ clones were selected that had 1) DNA Sanger sequencing predicted to cause loss of one Z18 allele and 2) which had 50% protein reduction by immunoblot. $Z18^{trunc/+}$ single cell clones were screened by Sanger sequencing and expression of the truncated protein isoform was confirmed by immunoblotting. Wild-type Z18 (hereafter $Z18^{WT}$) control cell lines were generated from cells transfected with Cas9-expressing vectors with no gRNA. These were then single-cell cloned and confirmed to have intact Z18 via Sanger sequencing and immunoblot.

*Bulk Z18 CRISPR:* Since Z18 null clones were not able to be isolated as expected (Z18 is a common essential gene in DepMap[18]), transient bulk CRISPR sequencing was executed using the most efficient guide (gRNA 1, Fig. S3A, Table S5) in triplicate. Cells transfected with a Cas9-expressing vector without gRNA served as controls. Transfected cells were selected with puromycin for 48 hours and Z18 protein reduction was confirmed by immunoblot.

*$Z18^{trunc}$-expressing lines:* pLENTI CMV Clover (fluorescent control) and pLENTI CMV $Z18^{trunc}$-expressing human melanoma A375 cell lines were generated via lentiviral transduction and stable puromycin selection. Lines were made in biologic triplicate and maintained in selection antibiotic.

**RNA-seq:** RNA was isolated and genomic DNA was removed from control (n=2), $Z18^{trunc/+}$ (n=3), and $Z18^{-/+}$ (n=3) single cell clones and from bulk-CRISPR control (n=3) and Z18-CRISPR (n=3) using the method as described above for zebrafish. All samples were polyA-selected (NEB, E7490) and underwent stranded library preparation (NEB, E7760S). Samples were sequenced on a HiSeq 4000 with paired end sequencing.

**RNA-seq analysis**: Data processing was performed using SQuIRE[10] as described above for zebrafish TE quantification, except data were mapped to hg38. GSEA[19] was done on RefSeq DESeq2[15] output from SQuIRE.

**qPCR:** RNA was isolated (Zymo Research, 2052). First-strand cDNA was synthesized (Takara Bio, RR036A) and used for qPCR (Biotium, 31003) analysis. RNA expression was plotted as compared to GAPDH. Primers used for qPCR are listed in Table S6.

**4sU pulse-chase RNA decay measurement:** Control Clover- and Z18$^{trunc}$-expressing cell lines were labeled with 50 μM 4-thiouridine (Sigma Aldrich, T4509) for 2 hours followed by replacement with 4sU-free media. Cells were harvested at 0 and 4 hours post media replacement. Total RNA was isolated using Trizol and treated with DNase I (Thermofisher, 18068015). 10 μg 4sU-labeled RNA was biotinylated, purified, captured by streptavidin magnetic beads (Invitrogen, 65002), eluted using 100 mM dithiothreitol (DTT), and purified by ethanol precipitation. cDNA was prepared and quantified by qPCR as above.

**Nascent RNA qPCR:** The TT-seq protocol[20] was utilized with a qPCR readout. Clover- and Z18$^{trunc}$-expressing cell lines were labeled with 500 μM 4-thiouridine (Sigma Aldrich, T4509) for 5 min, lysed using Trizol, then RNA was extracted. 50 μg of total RNA was DNase I treated (Thermofisher, 18068015) and fragmented to 1000-1200 base pair fragments with a Covaris E220 ultrasonicator (50 sec at 105 peak incident power). Nascent RNA was biotinylated, purified, captured by streptavidin magnetic beads (Invitrogen, 65002), eluted using 100 mM DTT, and purified by ethanol precipitation. cDNA was prepared and quantified by qPCR as above.

**Western blotting:** 5–15 μg of protein was run on 10% tris glycine gels followed by either semi-dry or wet (30V, overnight) transfer to PVDF or nitrocellulose membranes. Membranes were blocked with 5% skim milk in TBST. The following primary antibodies were used with 1:1000 dilution: anti-Z18 (Novus, NBP1-82570), anti-ZCCHC8 (Proteintech, 23374-1-AP), anti-PABPC1 (abcam, ab21060), anti-SRRT (Invitrogen, PA5-31593), anti-GAPDH (CST, 2118), anti-VCL (Sigma, HPA002131), and anti-Beta Actin (CST, 3700). These antibodies were used with a 1:2000 dilution: Anti-V5 (abcam, ab27671) and anti-MTR4 (abcam, ab70551). Anti-LINE1 ORF1p (EMD Millipore, clone 4H1) was used with 1:5000 dilution. Rabbit or mouse secondary antibodies were diluted to 1:2000 and incubated for 1 hour at room temperature (20-25°C) (HRP: CST, 7074S or 7076S) (fluorescent: Licor 926-32211 or 926-68070). When visualizing immunoprecipitated protein, TidyBlot HRP (Biorad, STAR209P) was used at a 1:400 dilution. Films were developed with ECL substrate (Thermofisher 32106 or Cytivia RPN2232).

**Immunoprecipitation mass spectrometry (IP-MS):** Coding sequences of full-length Z18 (hereafter $Z18^{fl}$) and patient truncated $Z18^{trunc}$ (as described above in Zebrafish cloning) were assembled using the Gateway cloning system into pcDNA3.2 C-terminal V5 tag destination vectors (ThermoFisher, 12489019). Each construct was transiently transfected into A375 human melanoma cells in triplicate (Thermofisher, L3000008). Nuclei were isolated and lysed 48 hours after transfection (ThermoFisher, 78833). Anti-V5 (Clone V5-10, V8012 Sigma) was conjugated to protein G beads (ThermoFisher, 10004D) and used to IP V5-tagged proteins in the presence or absence of RNase A/T1 (100 µg/mL ThermoFisher, EN0551). IPed proteins were eluted by boiling then submitted for mass spectrometry at the Taplin Mass Spectrometry Facility at Harvard University. Data were filtered for proteins with >4 total peptides in all replicates in the most permissive condition ($Z18^{fl}$ -RNAse) and >4.5 the average peptide number compared to a previous control IP (Clover-V5)[21] (n=240 proteins). The % total peptides of identified proteins vs. bait Z18 was calculated. To identify significant changes in direct interactors, a 2-sided t-test was done between +RNase $Z18^{fl}$ and $Z18^{trunc}$ conditions. Proteins that were significantly changed between +RNase $Z18^{fl}$ and $Z18^{trunc}$ ($P<0.05$) and had an average >5% of bait in at least one +RNase condition were plotted in Graphpad Prism (n=8).

**Z18 zinc-finger domain structural modeling:** The protein sequence for Z18 was downloaded from Uniprot (Sequence ID Q86VM9) and the zinc-finger domain spanning residues 219 to 245 was selected for homology modeling. The Z18 zinc finger three dimensional structure was modeled using the online Swiss-Model server[22] with the RBM22 protein from the human spliceosome Cryo-EM structure with RNA oligomer (PDB 6ID1)[23] as a template.

**Modified PAR-CLIP:** The aromatic zinc finger (ZnF) mutations (F226A, F227A, and W234A) were introduced into $Z18^{trunc}$ using in vitro mutagenesis (NEB, E0554S). A prior study of the CCCH domain in the structurally similar ZC3H12 protein identified the aromatic residue equivalent to F240 to be unimportant for RNA binding; thus, we left F240 intact[24]. We were uncertain whether F226 or F227 were important because they are both conserved, so we mutated both. V5-tagged Clover (control), $Z18^{fl}$, $Z18^{trunc}$, and $Z18^{trunc}$ ZnF aromatic mutant (hereafter $Z18^{truncZnFmut}$) constructs were transiently transfected into A375 melanoma cells (Thermofisher, L3000008). PAR-CLIP was performed as published[25] except protein size selection was not completed. At 24 hours post-transfection, cells were labeled with 100 mM 4sU for 16 hours, media was removed, and cells were covered with PBS for UV crosslinking (2 min, 316 nm, Spectrolinker XL-1500). The crosslinked cell pellet was lysed with RIPA buffer (50 mg cell pellet/mL RIPA buffer), vortexed, and incubated on ice for 20 min before brief sonication (Covaris E220 ultrasonicator, 1 min, PIP 140.0, Duty factor 7.0, CPB 300). Protein G Dynabeads (330 µL/IP, Invitrogen, 10004D) were conjugated to anti-V5 (66 µg/IP, Sigma, V8012). IP was done in triplicate overnight at 4°C. Beads were washed once with RIPA buffer and twice with IP wash buffer (150 mM NaCl, 50 mM Tris). On-bead Proteinase K digestion (10 µL/IP, NEB, P8107S) was performed at 50 $^{o}$C for 45 min with gentle mixing every 5 min. RNA was isolated via acid Phenol-Chloroform extraction, purified using NEBNext RNA sample purification beads (NEB, E6351S), ribodepleted (NEB, E7400L), and prepared for stranded Illumina sequencing (NEB, E7760S).

**PAR-CLIP data analysis:**

Reads were trimmed and quantified using SQuIRE[10] as explained above. TEs that were found to be significantly bound to bait protein (Z18$^{fl}$, Z18$^{trunc}$, or Z18$^{truncZnFmut}$ ) vs. control Clover were identified using DE-seq2[15].

_Identification of bound regions:_ Trimmed reads were mapped to the human genome build hg38 using bowtie (v 1.1.2) in paired-end mode with parameters, -v 2 -m 10 –best –strata[26]. To identify high confidence clusters of T to C transitions resulting from UV crosslinking of 4sU-labeled RNA to Z18, WavCluster version 2.36.0[27] was used. To identify a unified set of regions of interest for each condition, the WavCluster output cluster.bed files were merged for replicates of each condition (Clover, Z18$^{fl}$, Z18$^{trunc}$, or Z18$^{truncZnFmut}$).

_Cluster annotation:_ The combined cluster files for each condition were annotated using bedtools intersect -c option, for: a) LTRs and LINEs using UCSC RepeatMasker (downloaded 11/08/2023), b) exons from refGene.gtf (downloaded 11/08/2023), and c) introns. Introns were annotated using a .bed file including all disjoint segments of canonical introns[28,29] using hg38.ncbiRefSeq.gtf[30] (those referring to patches or haplotypes were removed). The number of clusters annotated as overlapping with LTRs, LINEs, introns, or exons were calculated and plotted for each condition. The average cluster size (base pairs) from each condition was similar (Z18$^{fl}$ 137.51; Z18$^{trunc}$ 132.06; Clover 129.55).

*Quantification of bound regions:* Trimmed reads were remapped to the human genome build hg38 using the splice-aware mapper STAR (v. 2.5.3a) in paired-end mode with parameters --outSAMtype BAM SortedByCoordinate --outWigStrand Stranded --outWigType bedGraph[7]. To identify reads that overlapped Z18$^{fl}$ bound regions from WavCluster, each bam was intersected with the merged Z18$^{fl}$ using bedtools -intersect. Stringtie v. 1.3.3b[31] was used to count expression from intersected bam files annotated for: a) LTRs and LINEs, b) exons, and c) introns as above. Normalized reads per transcript were used as input for DEseq2[15] using a size factor of 1 comparing Z18$^{fl}$ or Z18$^{trunc}$ vs. Clover (ctrl). Log2 fold change of Z18$^{fl}$ or Z18$^{trunc}$ vs. Clover for LTRs, LINEs, introns, or exons was plotted and a non-parametric two-sided Wilcoxon rank sum test was used to compare clusters annotated to contain LTRs, LINEs, or introns vs. exons.
*Quantification of nucleotide base composition:* Clusters base composition was calculated. The genomic co-ordinates for each cluster .bed file were used to calculate the coverage in the corresponding BAM files using mpileup from samtools[32].

#### Figure S1

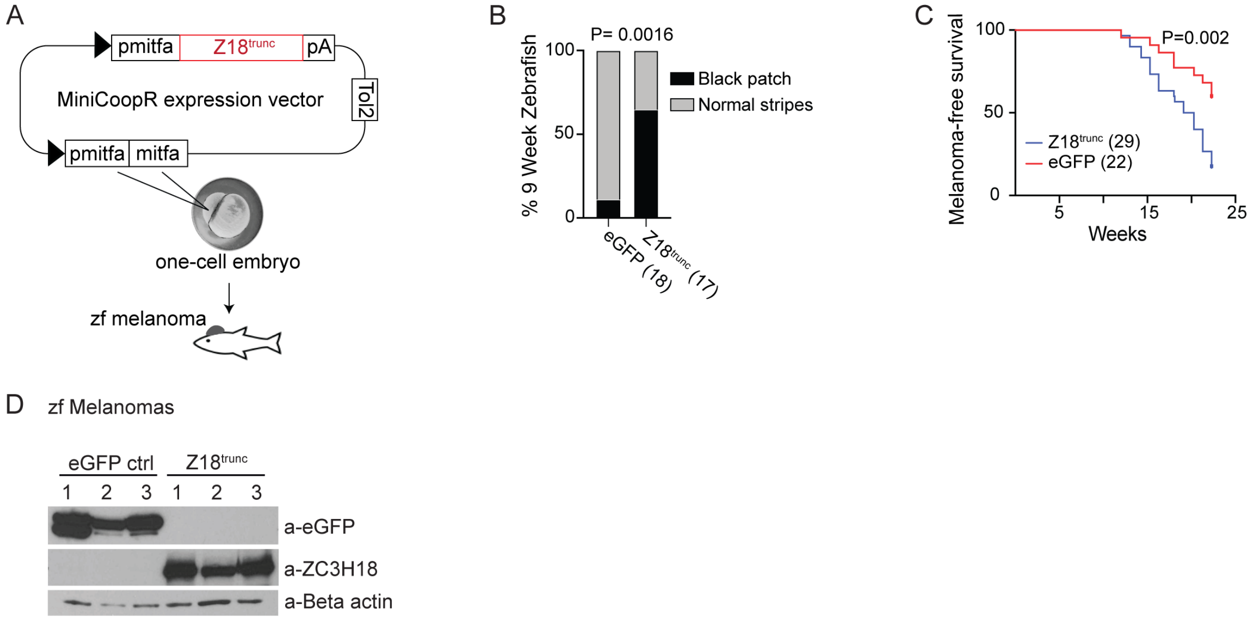


**Fig. S1. ZC3H18$^{trunc}$ Expression in Zebrafish Melanoma.** A) Schematic for Z18$^{trunc}$ expression in Triples using the MAZERATI system. pmitfa = zebrafish melanocyte promoter; Tol2 = sequences needed in cis for transgenesis. B) Quantification of black patches. P=0.0016 (Fisher's exact test). n=zebrafish. C) Percent melanoma-free survival. P=0.002 (log rank). n=zebrafish. D) Immunoblot for eGFP and human Z18$^{trunc}$ in eGFP- and Z18$^{trunc}$-expressing zebrafish melanomas.

#### Figure S2

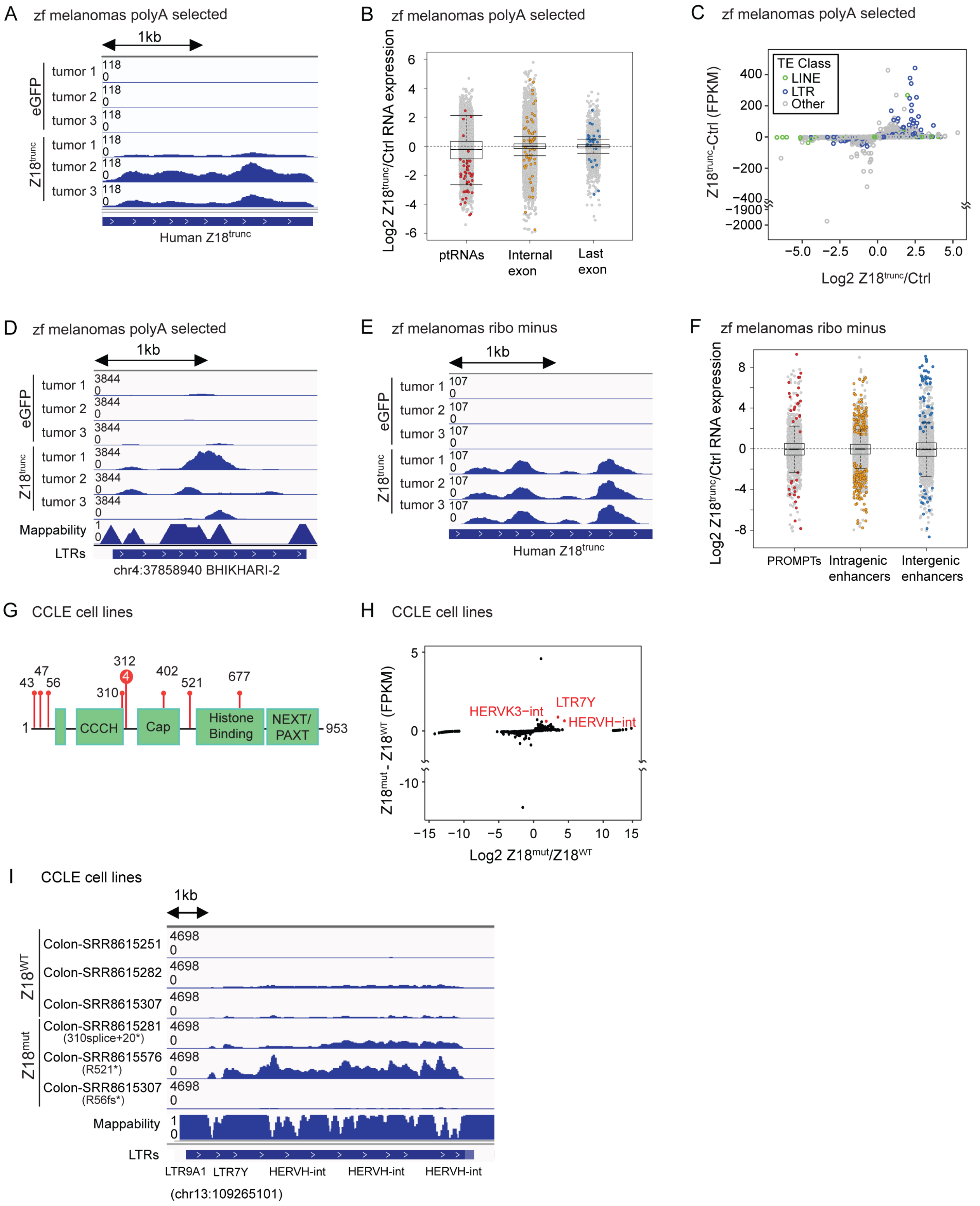


**Fig. S2. RNA-seq Defines ZC3H18$^{trunc}$ RNA Targets.** A) IGV plot showing human Z18$^{trunc}$ expression from pA-selected RNA-seq of eGFP- and Z18$^{trunc}$-expressing zebrafish melanomas. B)

Box plot showing log2 fold change of $Z18^{trunc}$ vs. control eGFP zebrafish melanomas for ptRNAs (left), internal exon (middle), or last exon (right). Color dots = significantly changed (q<0.1). ptRNA = RNA generated by usage of intronic polyadenylation sites. Internal exon = constitutive internal exon (non-alternatively spliced) isoform. C) Difference in average transposable element (TE) loci expression (FPKM) versus log2 fold change for $Z18^{trunc}$ (n=3) vs. eGFP (n=3) zebrafish melanomas. Circles = individual TE loci. D) IGV plot showing one example Bhikhari LTR (ERV) in eGFP- and $Z18^{trunc}$-expressing zebrafish melanomas. E) IGV plot showing human $Z18^{trunc}$ expression from ribo-minus RNA-seq of eGFP and $Z18^{trunc}$-expressing zebrafish melanomas. F) Box plot showing log2 fold change of $Z18^{trunc}$ vs. eGFP for promoter upstream transcripts (PROMPTs) (left), intragenic enhancers (middle), or intergenic enhancers (right). Color dots = significantly changed (q<0.1). G) Lollipop plot of Z18 mutations ($Z18^{mut}$) in CCLE cell lines (n=11). H) Difference in average expression (FPKM) vs. log2 fold change of significantly changed (Wilcoxon signed rank test, P<0.05) TE subfamilies from $Z18^{mut}$ vs. $Z18^{WT}$ CCLE cell lines. Subclasses with average difference in expression >0.5 and log2 fold change with >1.75 are labeled in red. Dots = subclasses. I) IGV plots of LTR (ERV) expression in $Z18^{mut}$ and disease-matched $Z18^{WT}$ CCLE cell lines. In box plots, the black horizontal line indicates the median, the box covers the interquartile range (IQR), and the whiskers extend to 1.5× the IQR.

### Figure S3

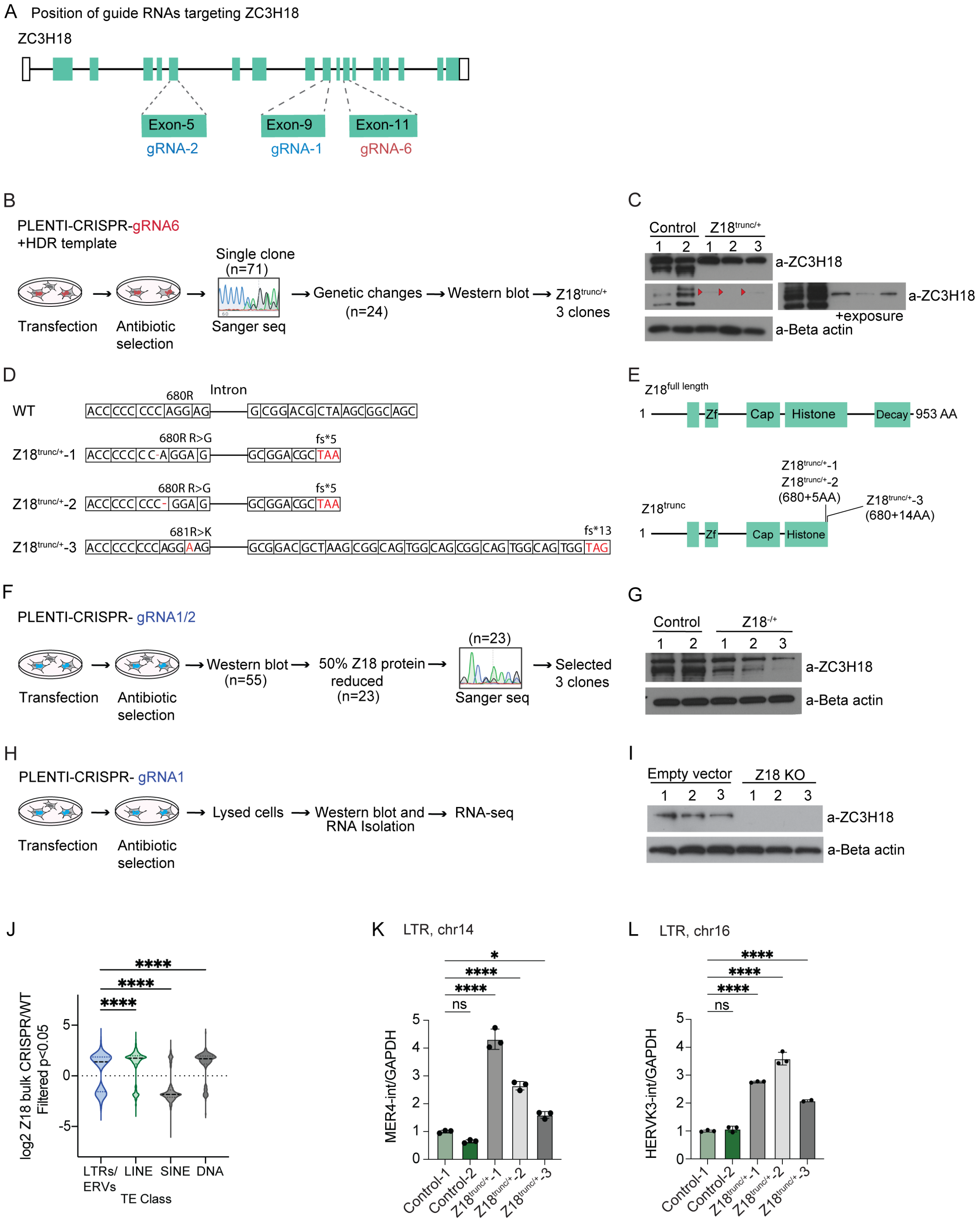
A
Position of guide RNAs targeting ZC3H18
ZC3H18
Exon-5
gRNA-2
Exon-9
gRNA-1
Exon-11
gRNA-6
B
PLENTI-CRISPR-gRNA6
+HDR template
Single clone
(n=71)
Transfection
Antibiotic selection
Sanger seq
Genetic changes
(n=24)
Western blot
Z18trunc/+
3 clones
C
Control
Z18trunc/+
1 2 1 2 3
a-ZC3H18
a-ZC3H18
+exposure
a-Beta actin
D
WT
680R
Intron
ACC CCC CCC AGG AG
G CGG ACG CTA AGC GGC AGC
Z18trunc/+-1
680R R>G
fs*5
ACC CCC C C-A GGA G
GC GGA CGC TAA
Z18trunc/+-2
680R R>G
fs*5
ACC CCC CCC - GGA G
GC GGA CGC TAA
Z18trunc/+-3
681R>K
fs*13
ACC CCC CCC AGG AAG
GCG GAC GCT AAG CGG CAG TGG CAG CGG CAG TGG CAG TGG TAG
E
Z18full length
1
Zf
Cap
Histone
Decay
953 AA
Z18trunc/+-1
Z18trunc/+-2
(680+5AA)
Z18trunc/+-3
(680+14AA)
Z18trunc
1
Zf
Cap
Histone
F
PLENTI-CRISPR- gRNA1/2
Transfection
Antibiotic selection
Western blot
(n=55)
50% Z18 protein reduced
(n=23)
(n=23)
Sanger seq
Selected 3 clones
G
Control
Z18-/+
1 2 1 2 3
a-ZC3H18
a-Beta actin
H
PLENTI-CRISPR- gRNA1
Transfection
Antibiotic selection
Lysed cells
Western blot and RNA Isolation
RNA-seq
I
Empty vector
Z18 KO
1 2 3 1 2 3
a-ZC3H18
a-Beta actin
J
log2 Z18 bulk CRISPR/WT
Filtered p<0.05
5
0
-5
****
****
****
LTRs/ ERVs
LINE
SINE
DNA
TE Class
K
LTR, chr14
MER4-int/GAPDH
*
****
****
ns
0
1
2
3
4
5
Control-1
Control-2
Z18trunc/+-1
Z18trunc/+-2
Z18trunc/+-3
L
LTR, chr16
HERVK3-int/GAPDH
****
****
****
ns
0
1
2
3
4
5
Control-1
Control-2
Z18trunc/+-1
Z18trunc/+-2
Z18trunc/+-3

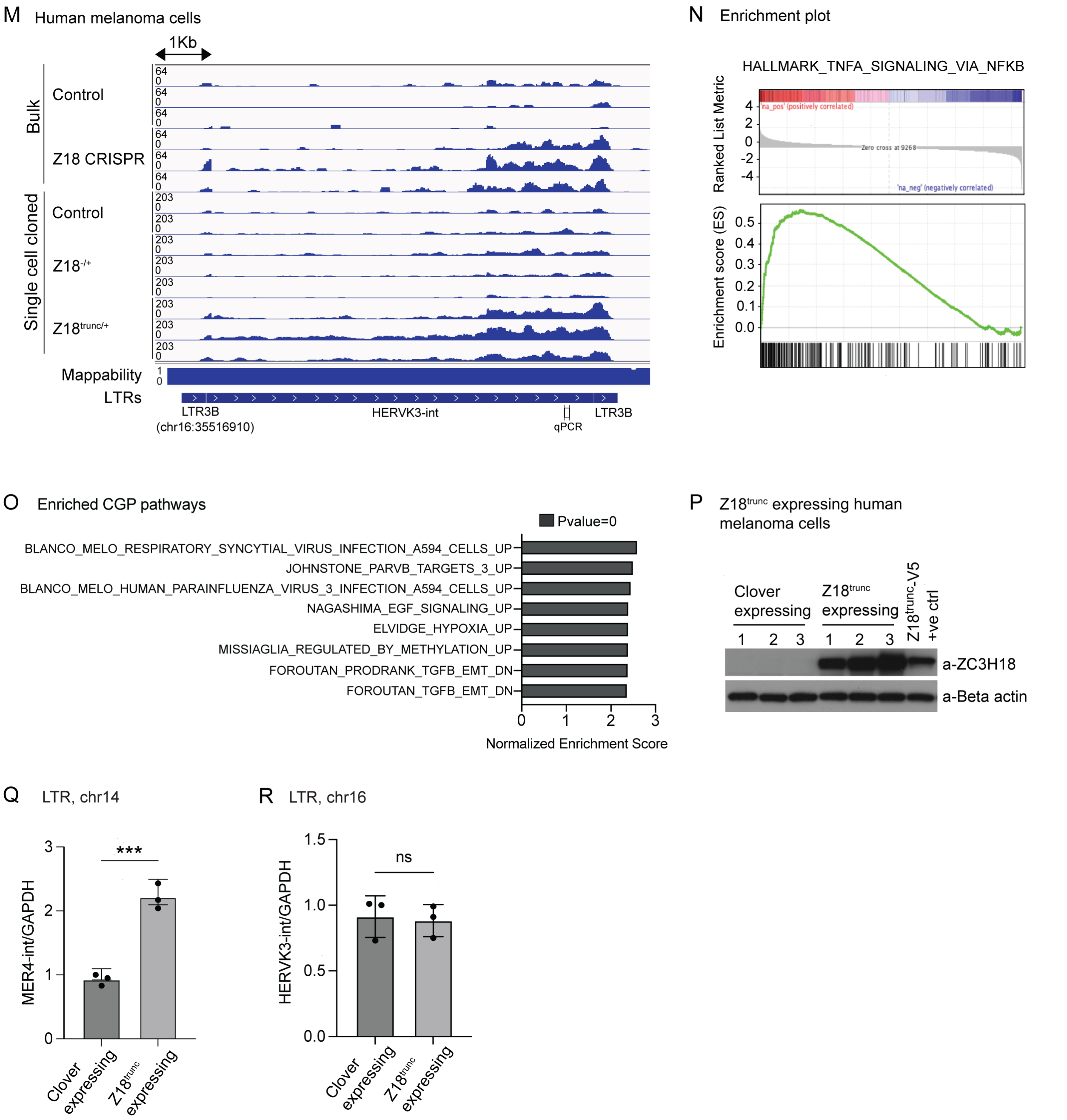


**Fig. S3. Validation of ZC3H18 Mutations in Human Melanoma Cell Lines.** A) Schematic showing sgRNA target positions on the *Z18* gene. B) Workflow for generation of heterozygous $Z18^{trunc/+}$ cell lines. C) Z18 immunoblot of $Z18^{WT}$ control and $Z18^{trunc/+}$ cell lines. Red arrows = new truncated isoform. Longer exposure on right. D-E) Diagram showing the effects of $Z18^{trunc}$ mutations in the selected cell lines vs. intact Z18 on D) RNA and E) protein. F) Workflow and G) immunoblot for $Z18^{WT}$ control and heterozygous loss-of-function $Z18^{-/+}$ cell lines. H) Workflow and I) immunoblot of control vs. bulk Z18 CRISPR. J) Log2 fold change expression of significantly changed TEs ($P<0.05$) from bulk Z18 CRISPR (n=3) vs. control (n=3). Thick dotted line = median. Light dotted line = quartiles. **** $P<0.0001$ (Ordinary one-way ANOVA). K-L) qPCR for two LTRs (ERVs) vs. GAPDH from control (n=2) and $Z18^{trunc/+}$ (n=3) cell lines. Mean ± SD. * P=0.01, **** $P<0.0001$ (Ordinary one-way ANOVA). M) IGV plot of chr16 HERVK3-int ERV expression in bulk Z18 CRISPR vs. control and $Z18^{-/+}$, $Z18^{trunc/+}$, and $Z18^{WT}$ cell lines. N-O) GSEA for $Z18^{trunc/+}$ (n=3) vs. $Z18^{WT}$ control (n=2) cell line RNA-seq. N) Enrichment plot for the most significantly changed Hallmark pathway. O) Normalized enrichment scores for the most significantly changed Chemical and Genetic Perturbation (CGP) pathways. P) Z18 immunoblot of Clover and $Z18^{trunc}$-expressing human melanoma cell lines. Q-R) qPCR analysis of Q) chr14 MER4-int as defined in Fig. 3E and R) chr16 HERVK3-int as defined in Fig. S3M in Clover- (n=3) and $Z18^{trunc}$-expressing (n=3) cell lines. Mean ± SD. *** P=0.0005, ns=non-significant (Unpaired t test, two-sided).

#### Figure S4

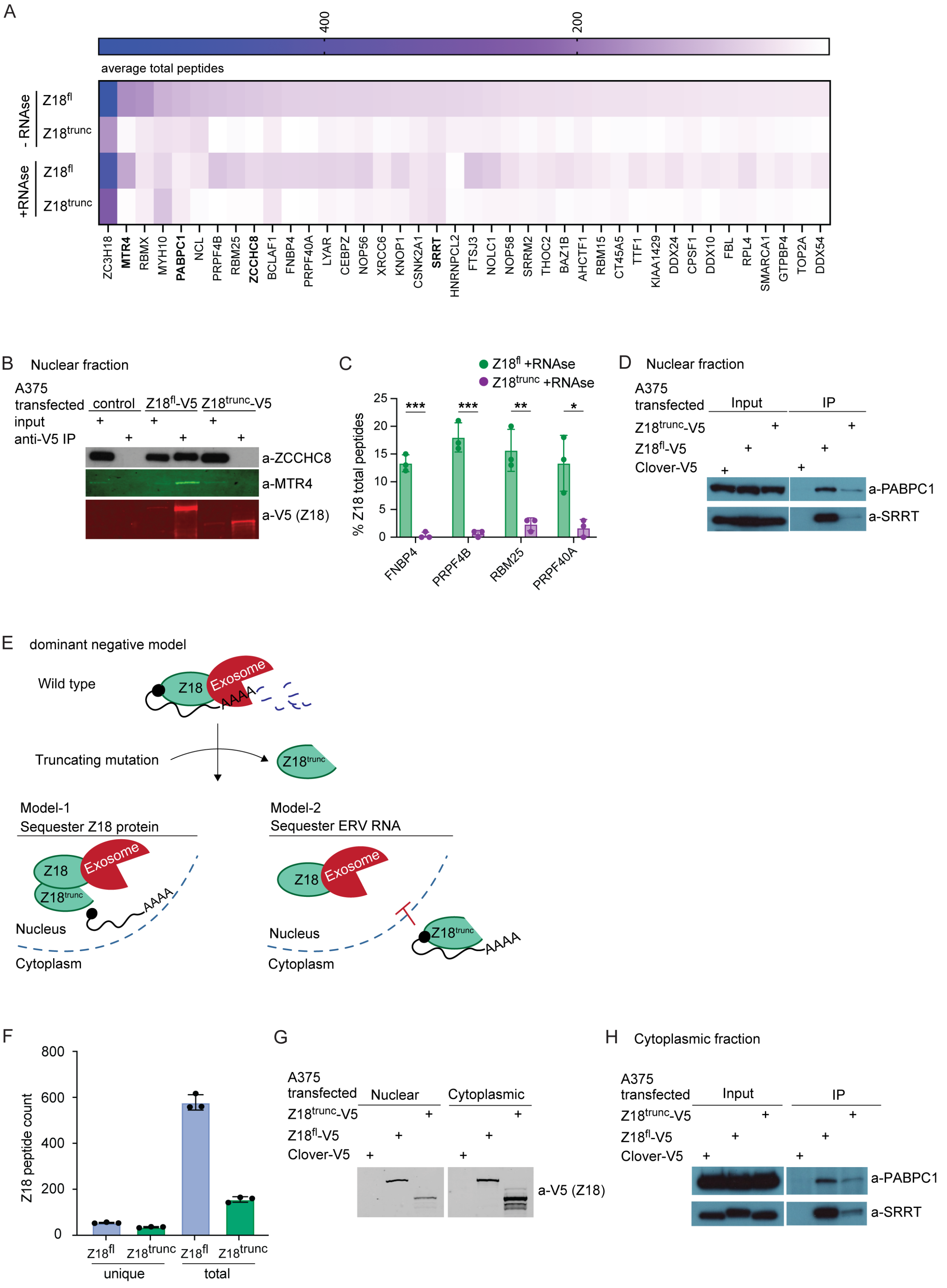
A
400
200
average total peptides
- RNAse
+RNAse
Z18fl
Z18trunc
ZC3H18
MTR4
RBMX
MYH10
PABPC1
NCL
PRPF4B
RBM25
ZCCHC8
BCLAF1
FNBP4
PRPF40A
LYAR
CEBPZ
NOP56
XRCC6
KNOP1
CSNK2A1
SRRT
HNRNPCL2
FTSJ3
NOLC1
NOP58
SRRM2
THOC2
BAZ1B
AHCTF1
RBM15
CT45A5
TTF1
KIAA1429
DDX24
CPSF1
DDX10
FBL
RPL4
SMARCA1
GTPBP4
TOP2A
DDX54
B Nuclear fraction
A375
transfected
control
Z18fl-V5
Z18trunc-V5
input
anti-V5 IP
a-ZCCHC8
a-MTR4
a-V5 (Z18)
C
Z18fl +RNAse
Z18trunc +RNAse
% Z18 total peptides
FNBP4
PRPF4B
RBM25
PRPF40A
D Nuclear fraction
A375
transfected
Input
IP
Z18trunc-V5
Z18fl-V5
Clover-V5
a-PABPC1
a-SRRT
E dominant negative model
Wild type
Z18
Exosome
AAAA
Truncating mutation
Z18trunc
Model-1
Sequester Z18 protein
Nucleus
Cytoplasm
Model-2
Sequester ERV RNA
F
Z18 peptide count
unique
total
G
A375
transfected
Nuclear
Cytoplasmic
Z18trunc-V5
Z18fl-V5
Clover-V5
a-V5 (Z18)
H Cytoplasmic fraction
A375
transfected
Input
IP
a-PABPC1
a-SRRT

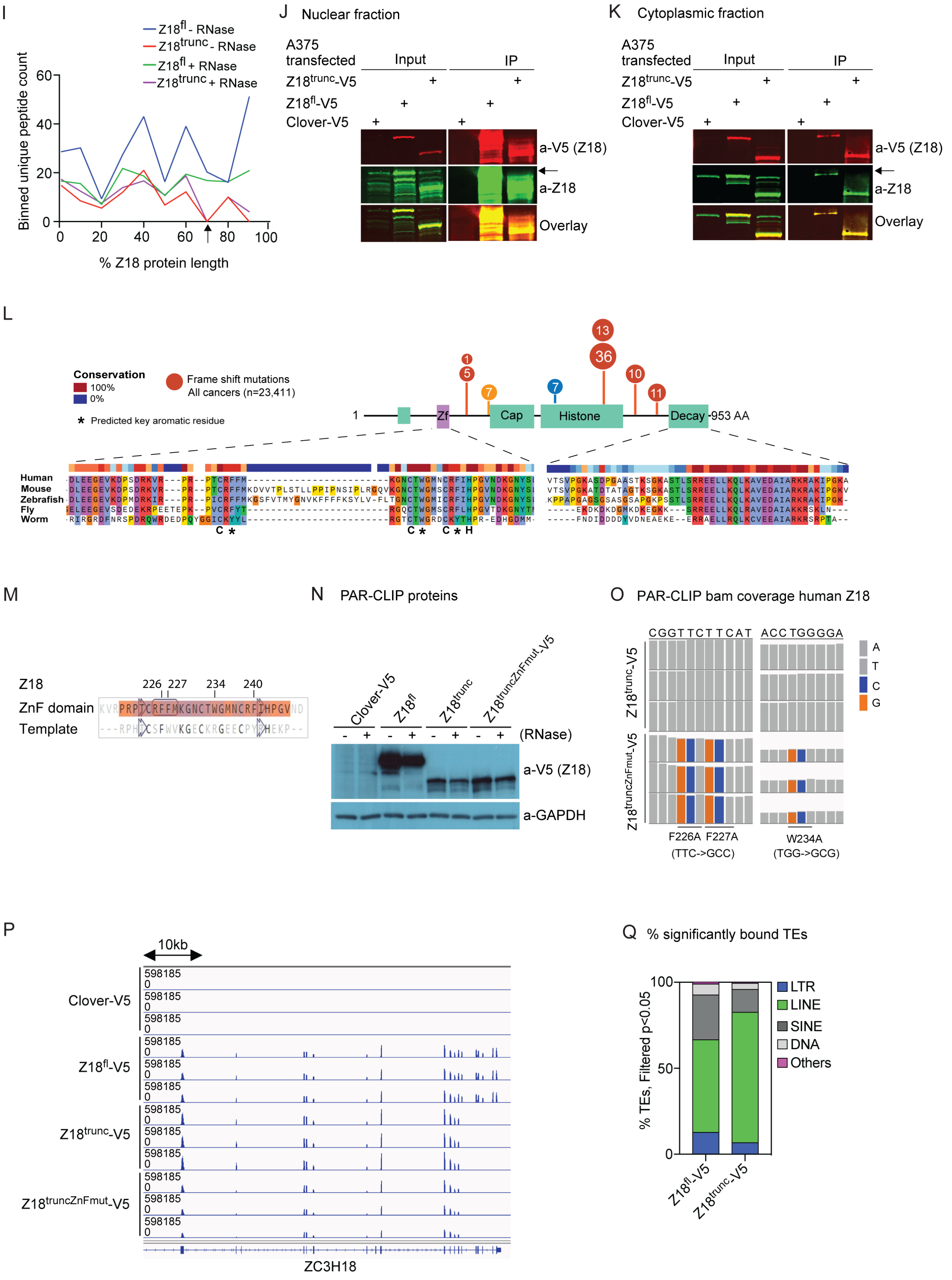
I
Z18fl - RNase
Z18trunc - RNase
Z18fl + RNase
Z18trunc + RNase
Binned unique peptide count
% Z18 protein length
J Nuclear fraction
A375 transfected
Input
IP
Z18trunc-V5
Z18fl-V5
Clover-V5
a-V5 (Z18)
a-Z18
Overlay
K Cytoplasmic fraction
L
Conservation
100%
0%
Frame shift mutations
All cancers (n=23,411)
Predicted key aromatic residue
1
Zf
Cap
Histone
Decay
953 AA
Human
Mouse
Zebrafish
Fly
Worm
M
Z18
ZnF domain
Template
226 227 234 240
N PAR-CLIP proteins
Clover-V5
Z18fl
Z18trunc
Z18truncZnFmut-V5
(RNase)
a-V5 (Z18)
a-GAPDH
O PAR-CLIP bam coverage human Z18
CGGTTCTTCAT
ACCTGGGGA
Z18trunc-V5
Z18truncZnFmut-V5
A
T
C
G
F226A F227A
(TTC->GCC)
W234A
(TGG->GCG)
P
10kb
Clover-V5
Z18fl-V5
Z18trunc-V5
Z18truncZnFmut-V5
598185
ZC3H18
Q % significantly bound TEs
% TEs, Filtered p<0.05
LTR
LINE
SINE
DNA
Others
Z18fl-V5
Z18trunc-V5

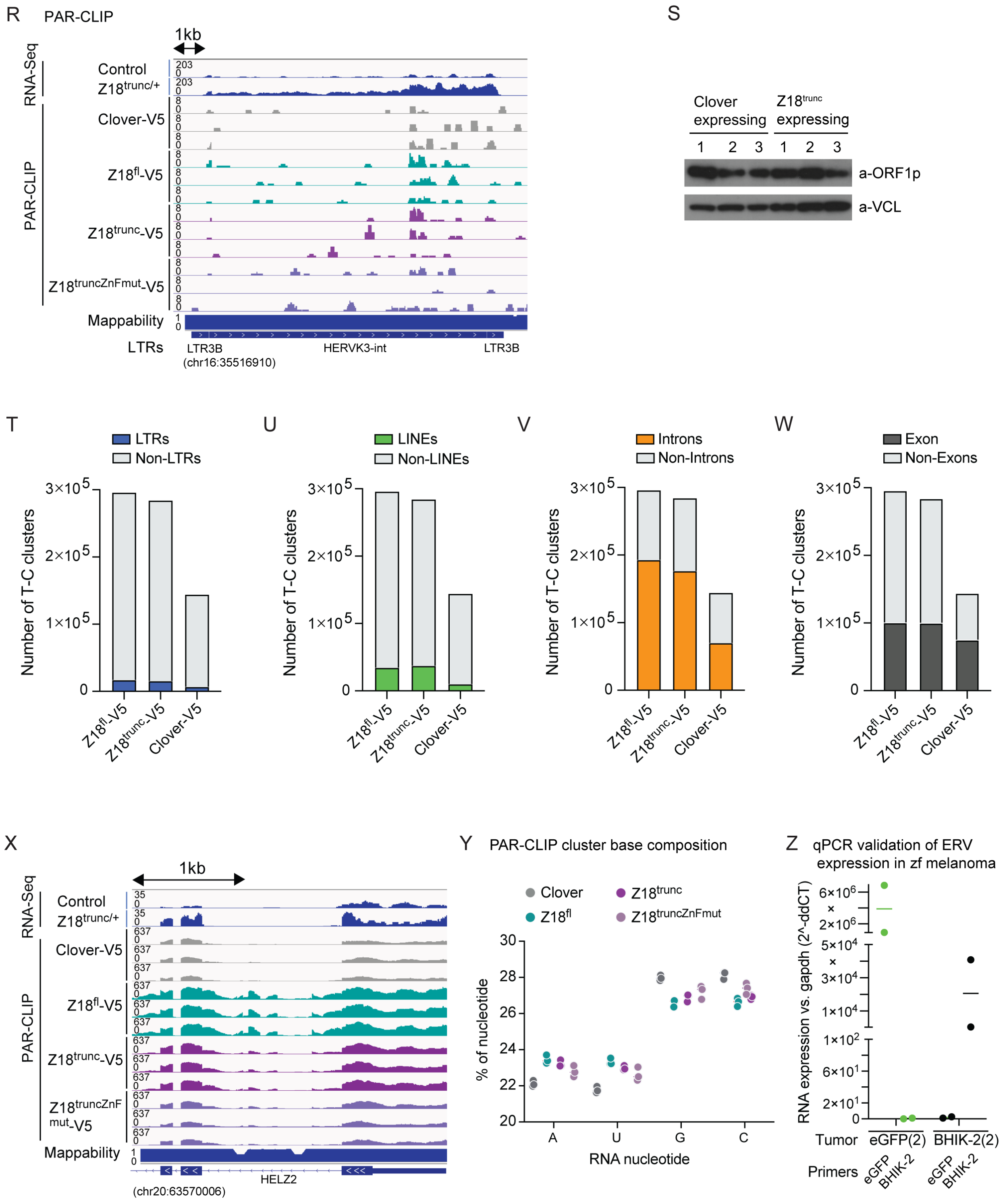
R PAR-CLIP
1kb
RNA-Seq
Control
Z18trunc/+
PAR-CLIP
Clover-V5
Z18fl-V5
Z18trunc-V5
Z18truncZnFmut-V5
Mappability
LTRs
LTR3B
(chr16:35516910)
HERVK3-int
LTR3B
S
Clover expressing
Z18trunc expressing
1 2 3 1 2 3
a-ORF1p
a-VCL
T
LTRs
Non-LTRs
Number of T-C clusters
Z18fl-V5
Z18trunc-V5
Clover-V5
U
LINEs
Non-LINEs
V
Introns
Non-Introns
W
Exon
Non-Exons
X
1kb
RNA-Seq
Control
Z18trunc/+
PAR-CLIP
Clover-V5
Z18fl-V5
Z18trunc-V5
Z18truncZnFmut-V5
Mappability
HELZ2
(chr20:63570006)
Y PAR-CLIP cluster base composition
Clover
Z18fl
Z18trunc
Z18truncZnFmut
% of nucleotide
RNA nucleotide
A U G C
Z qPCR validation of ERV expression in zf melanoma
RNA expression vs. gapdh (2^-ddCT)
Tumor
eGFP(2)
BHIK-2(2)
Primers
eGFP
BHIK-2

**Fig. S4. Dominant Negative Model for ZC3H18$^{trunc}$.** A) Heat map of average total peptides from immunoprecipitation mass spectrometry (IP-MS) of V5-tagged Z18$^{fl}$ and Z18$^{trunc}$ +/- RNAse. n=3 for all conditions. Visualized proteins had total peptides >4.5× control IP and >4 peptides in each replicate. Top 40 of 240 proteins shown. B) V5 IP from nuclear extract and immunoblot for ZCCHC8, MTR4, and V5 from cells expressing V5-tagged Clover, Z18$^{fl}$, or Z18$^{trunc}$. C) Total peptide counts normalized to Z18 bait for other four significantly differentially bound proteins in Z18$^{fl}$ vs. Z18$^{trunc}$. Mean ± SD. FNBP4 *** P=0.0002, PRPF4B *** P=0.00034, RBM25 ** P=0.0032, PRPF40A * P=0.015 (2-sided t-test). n=3 for all conditions. D) V5 IP from nuclear extract and immunoblot detecting PABPC1 and SRRT from cells expressing V5-tagged Clover, Z18$^{fl}$, or Z18$^{trunc}$. E) Schematic depicting two models of Z18$^{trunc}$ dominant negative activity. F) IP-MS unique and total Z18 peptide counts. Mean ± SD. n=3 for all conditions. G) V5 IP from nuclear or cytoplasmic fractions and V5 immunoblot from cells expressing V5-tagged Clover, Z18$^{fl}$, or Z18$^{trunc}$. H) V5 IP from cytoplasmic extract and immunoblot detecting PABPC1 and SRRT from cells expressing V5-tagged Clover, Z18$^{fl}$, or Z18$^{trunc}$. I) IP-MS binned total average Z18 peptide counts plotted by % Z18 protein length. Average from n=3 for all conditions. J-K) V5 IP and immunoblot for V5 and Z18 from cells expressing V5-tagged Clover, Z18$^{fl}$, or Z18$^{trunc}$. J) Nuclear protein and K) cytoplasmic protein. Arrows highlight that Z18$^{trunc}$ does not IP endogenous Z18$^{fl}$. L) Protein alignment showing conservation of the Z18 zinc-finger and RNA surveillance recruitment domains across 5 species. C-C-C-H = residues predicted to coordinate with zinc to form the zinc-finger domain structure. * = aromatic residues predicted to be involved in RNA coordination. M) Target-template protein sequence alignment for Z18 homology modeling. Top: Z18 zinc-finger domain. Bottom: RBM22. N) Immunoblot for V5 or GAPDH from PAR-CLIP input protein from cells expressing V5-tagged Clover, Z18$^{fl}$, Z18$^{trunc}$, or Z18$^{trunc}$ ZnF aromatic mutant (Z18$^{truncZnFmut}$) after 4sU labeling and crosslinking. O-P) IGV plot from PAR-CLIP RNA-seq showing O) Z18$^{truncZnFmut}$ (F226A, F227A, and W234A) mutations vs. Z18$^{trunc}$ and P) verifying length of Z18$^{fl}$, Z18$^{trunc}$, or Z18$^{truncZnFmut}$ transgene expression. n=3 replicates for all conditions. Q) Percent significantly bound (P<0.05) TEs from Z18$^{fl}$-V5 or Z18$^{trunc}$-V5 vs. Clover from PAR-CLIP SQuIRE analysis. R) IGV plot showing binding of Z18$^{fl}$, Z18$^{trunc}$, Z18$^{truncZnFmut}$, and Clover to HERVK3-int LTR/ERV. S) LINE1 ORF1p immunoblot from Clover- and Z18$^{trunc}$-expressing human melanoma cell lines. n=independently derived cell lines. T-W) Merged clusters of T-to-C transitions from Z18$^{fl}$-V5, Z18$^{trunc}$-V5, and Clover-V5 annotated for overlap with T) LTRs, U)

LINEs, V) introns, and W) exons. Clover clusters represent non-specific RNA binding (background). X) IGV plot of Z18-bound intron-containing transcript. Y) Nucleotide quantification in T-To-C transition clusters. Clover represents base composition of non-specific RNA binding (background). n=3 PAR-CLIP replicates. Mean ± SD. Z) qPCR of eGFP and BHIK-2 from eGFP-expressing control and BHIK-2-expressing zebrafish melanomas. BHIK-2=BHIKHARI-2. n=melanomas.

**Table S1. MSI status of tumors with Z18 mutations from TCGA.**

| MSI score | Z18 mutations | $Z18^{WT}$ |
|---|---|---|
| MSI high (>10) | 75 | 244 |
| MSI not high (<10) | 137 | 10326 |

**Table S2. Average expression and statistics of major TE classes from zebrafish pA-selected RNA-seq.** Mean expression of individual TE loci across TE classes for control eGFP (n=3) and $Z18^{trunc}$ (n=3). TE elements are ranked by the average expression difference between $Z18^{trunc}$ and eGFP.

| CLASS | MEAN EXPRESSION (eGFP) | MEAN EXPRESSION ($Z18^{trunc}$) | WILCOXON SIGNED-RANK TEST P-VALUE | WELCH'S T-TEST P-VALUE |
|---|---|---|---|---|
| LTR | 0.3715657 | 0.55457882 | 4.54e-12 | 0.002693 |
| SINE | 0.34140055 | 0.38899776 | 1.34e-14 | 0.5379 |
| LINE | 0.21240962 | 0.23041408 | 0.0008406 | 0.5006 |
| DNA | 0.40962196 | 0.40551719 | <2.2e-16 | 0.8143 |

**Table S3. Z18 mutations from CCLE.**

| CCLE cell lines with Z18 mutations | | | | | | |
|---|---|---|---|---|---|---|
| Cancer Type | Gender | Patient ID | Z18 gene Mutations | Z18 Protein Changes | Z18 protein Truncation | Genetically Altered Oncogene/Tumor Suppressor |
| Endometrial /Uterine Cancer | Female | SRR8615235 | c.2029_2030insCC | p.T677fs | Truncating (fs+9*) | TP53, PTEN, PIK3R1, ARID1A |
| | Female | SRR8615743 | c.(934-936)ttafs | p.L312fs | Truncating (fs+21*) | TP53, PTEN, PIK3R1, ARID1A, MSH2, MSH6 |
| | Female | SRR8615277 | Cell line1_c.935_936insA | p.LK312fs | Truncating (fs+13*) | TP53, PTEN, PIK3R1, ARID1A, POLE, CTNNB1, MSH2, MSH6 |
| | Female | SRR8615553 | Cell line2_c.935_936insA | p.LK312fs | Truncating (fs+13*) | CTNNB1, PIK3CA, PIK3R1, ARID1A, MSH6 |
| Colon/Colorectal Cancer | Male | SRR8615576 | Cell line1_c.1561C>T | p.R521* | Truncating (521*) | BRAF, MLH1, MSH6, APC, TP53, MUTYH |
| | Male | SRR8615281 | intron.929T>C | p.310splice+20* | Truncating (310+20*) | KRAS, MLH1, MSH6, APC, TP53, |
| | Male | SRR8615906 | c.167delG | p.R56fs | No protein | BRAF, MSH2, MSH6, APC, TP53 |
| Lung Cancer | Male | SRR8615600 | c.1204C>T | p.R402* | Truncating (402*) | KRAS |
| | Male | SRR8615988 | c.126_127insTT | p.V43fs | | BRAF |
| Gastric Cancer | Female | SRR8615672 | c.(934-936)ttafs | p.L312fs | Truncating (fs+21*) | MSH2, TP53, PIK3CA, KRAS, APC |
| Lymphoma | Male | SRR8616119 | c.139_146delGATCTGGA | p.DLE47fs | No protein | TP53, ARID1A, SETDB1 |

| CCLE cancer-type-matched cell lines with intact Z18 | | | | |
|---|---|---|---|---|
| Cancer Type | Gender | Patient ID | Z18 gene | Genetically Altered Oncogene/Tumor Suppressor |
| Endometrial /Uterine Cancer | Female | SRR8615236 | WT | MSH6, TP53, PTEN, PIK3CA, ARID1A |
| | Female | SRR8615237 | WT | TP53, PIK3CA |
| | Female | SRR8615275 | WT | MSH6, PMS2, TP53, PIK3CA, ARID1A, KRAS, CTNNB1 |
| | Female | SRR8615276 | WT | MLH1, MSH6, PTEN, PIK3CA, ARID1A, POLE |
| Colon/Colorectal Cancer | Female | SRR8615251 | WT | BRAF, MLH1, MSH2, APC, TP53 |
| | Male | SRR8615282 | WT | KRAS, MLH1, MSH6 |
| | Female | SRR8615307 | WT | APC |
| Lung Cancer | Male | SRR8615245 | WT | BRAF, TP53 |
| | Female | SRR8615252 | WT | TP53 |
| Gastric Cancer | Female | SRR8615225 | WT | ARID1A |
| Lymphoma | Male | SRR8615263 | WT | TP53 |

**Table S4. Median Log2FC of significantly changed TEs (p<0.05) from major TE classes in Z18-mutated vs. control CCLE cell lines.**

| TE class | Median Log2FC |
|---|---|
| LTR | 0.8758 |
| LINE | 0.8623 |
| SINE | 0.7370 |
| DNA | 0.8417 |

**Table S5. Cas9-sgRNA used for Z18 CRISPR.**

| | Sequence 5′-3′ (underlined=NGG) | Strand | Z18 Exon |
|---|---|---|---|
| **gRNA1** | ATGAAGACCGGGATCGCGAC<u>CGG</u> | - | 9 |
| **gRNA2** | TGACCATTGGCGAAGACGAA<u>CGG</u> | + | 5 |
| **gRNA6** | CGCCGGAGTGCTCACCTCCT<u>GGG</u> | - | 11 |

**Table S6. qPCR primers.**

| Primers | | 5′-3′ sequence |
|---|---|---|
| chr14 MER4_int_ | Fw | CTTGGGGTCTTGATCGGGAC |
| chr14 MER4_int | Rev | ATCTGCCTTCCACCGTCATG |
| chr16 HERVK3_int | Fw | TCAGCTTACTCATGGCACTCGT |
| chr16 HERVK3_int | Rev | TGAGGACACCACAACAGAGGG |
| GAPDH | Fw | CAAGAGCACAAGAGGAAGAGAG |
| GAPDH | Rev | CTACATGGCAACTGTGAGGAG |
| BHIKHARI-2 | Fw | CTTCTAAGTCATTAATTCCTGCAGCCCGGG |
| BHIKHARI-2 | Rev | TAACCCTCACTAAAGGGAACAAAAGCTGG |
| eGFP | Fw | GCGAGGGCGATGCCACCTACG |
| eGFP | Rev | TCACCTCGGCGCGGGTCTTGT |

**Table S7. Statistics for PAR-CLIP from Figure 4H-I comparing binding of LTRs, LINEs, and introns to exons (medians and p-values from Wilcoxon rank sum).**

| | **Medians** | | | |
|---|---|---|---|---|
| | LTRs | LINEs | Introns | Exons |
| Z18$^{fl}$ vs. Clover | 1.38 | 1.86 | 1.78 | 0.73 |
| Z18$^{trunc}$ vs. Clover | 0.96 | 1.48 | 1.22 | 0.53 |

| | **P values compared to exons** | | |
|---|---|---|---|
| | LTR | LINE | Introns |
| Z18$^{fl}$ vs. Clover | 9.92E-179 | 0.00E+00 | 0.00E+00 |
| Z18$^{trunc}$ vs. Clover | 3.03E-100 | 0.00E+00 | 0.00E+00 |

**Data S1. Z18 mutations in cancer.**
**Data S2. GSEA analysis from Z18$^{trunc/+}$ vs. control pA-selected RNA-seq.**
**Data S3. Z18 interacting partners identified from IP-MS analysis.**
